# Leupaxin inhibits the durotaxis and mechanosensitivity of metastatic breast cancer cells

**DOI:** 10.64898/2026.02.03.703253

**Authors:** Mathilde Mathieu, Jouni Härkönen, Monika Vaitkevičiūtė, Hellyeh Hamidi, Johanna Ivaska

## Abstract

Efficient cancer cell migration is essential for invasion and metastasis and is driven by cancer cell interaction with their extracellular matrix (ECM). Thus, ECM properties determine the migration phenotype. For example, ECM stiffness can guide cancer cell migration through durotaxis; however, the mechanisms regulating cancer cell durotaxis remain poorly understood. Using in-house stiffness gradient hydrogels, we discovered that MDA-MB-231 breast cancer cell metastatic variants selected for either bone or brain organotropism display impaired durotaxis. Moreover, akin to cells conditioned on soft substrates, these cells have altered mechanoresponses to increasing stiffness, including reduced cell spreading and focal adhesion assembly and signaling. We observed upregulation of the LIM-domain protein leupaxin, a focal adhesion component, in these variants and in soft-conditioned MDA-MB-231 cells compared to parental cells. Leupaxin silencing in brain-tropic metastatic cells restored durotaxis, whereas its overexpression in parental cells impaired durotaxis and mechanosensing, phenocopying the metastatic variants. Mechanistically, leupaxin disrupted paxillin–focal adhesion kinase (FAK) signaling, thereby inhibiting durotaxis. Together, these findings identify leupaxin as a negative regulator of durotaxis in metastatic cells and suggest a mechanism by which tumor cells adapt to mechanical heterogeneity to facilitate metastatic progression.

## Introduction

While migrating in tissues, cells encounter different environments with unique biomechanical properties, including stiffness, that impact cell behavior. In the main, stiffness-directed cell migration has been described as positive durotaxis, a preference to move from soft to stiffer matrices^1^. Most cell types undergo positive durotaxis in vitro^2–6^, including various cancer cell lines^7^. Moreover, positive durotaxis was recently pinpointed as a driver of metastatic pancreatic cancer progression^8^. However, the contribution of positive durotaxis in the progression of other cancers remains poorly understood or unexplored. In breast cancer, increased tumor rigidity has been generally associated with disease progression and increased cell proliferation^9–12^. However, as with most tumors, breast tumors are heterogenous in their physical properties, especially at the interface with the normal tissue; breast tumors exhibit a stiffer edge compared to the tumor core, and are surrounded by a softer healthy tissue^9,13^. Hence, for breast cancer to become metastatic, some tumor cells must be able to migrate away from the stiffer tumor microenvironment and invade into the softer healthy tissue. This clinically relevant in vivo scenario is in conflict with the positive durotaxis model. Instead, this “migration paradox”^14^ could be explained by the existence of a subset of tumor cells displaying either “negative” durotaxis, preferential migration from a stiff to a softer environment, or adurotaxis, stiffness-insensitive migration. We recently showed that glioblastoma U251-MG cells undergo negative durotaxis on synthetic stiffness-gradient gels^15^. Negative durotaxis has also been observed in an acral melanoma cell line^16^. In some cancer cell lines, such as the breast cancer MDA-MB-231 cell line, weakly adherent subpopulations have been described to have an adurotactic phenotype^17^. However, the molecular mechanism regulating these different durotactic behaviors remain unknown.

In addition to the initial invasion of surrounding tissue, cancer cells must adapt to alternating stiffnesses when colonizing secondary sites. Breast cancer metastasizes to the brain, bones, lungs and the liver^18^, which differs in their physical properties from the primary tumor^13,19–21^. The mechanisms driving metastatic cells’ organotropism remain mostly unknown and the contribution of durotaxis and mechanosensing have not been investigated. A well-established model of tissue-specific breast cancer metastasis are MDA-MB-231 variants selected by the Nishimura and Massagué labs for their ability to home preferentially to the bone or the brain^22,23^ (referred to as MDA-MB-231-Bo and MDA-MB-231-Br, or -Bo and -Br, respectively). Transcriptomic and proteomic profiling of these cells has led to the identification of genes enhancing their metastatic potential^24–26^, but their biomechanical properties and their putative role in their metastatic traits are not known.

To study the durotactic behavior of metastatic breast cancer cells, we used the MDA-MB-231-Br and -Bo cell lines. In addition, in an orthogonal approach, we conditioned parental MDA-MB-231 cells to different stiffness culture conditions. We discovered that the MDA-MB-231-Br and -Bo variants display attenuated durotaxis, mechanosensitivity and focal adhesion (FA) signaling, similarly to cells conditioned on soft substrates. Single-cell RNA sequencing revealed that elevated leupaxin expression is a shared feature of the -Br and -Bo variants when compared to parental cells. Furthermore, Leupaxin protein levels were higher in the metastatic variants and in the soft-conditioned cells compared to parental cells. We show that manipulating leupaxin levels (silencing or overexpression) determines the range of positive durotaxis and mechanosensitive signaling to YAP. We find that high leupaxin limits these events by inhibiting the paxillin-FAK pathway, identifying a previously unknown negative regulator of durotaxis.

### MDA-MB-231 brain and bone metastatic variants have reduced durotaxis and mechanosensitivity

Metastatic cells are able to colonize environments with different mechanical properties than the primary metastasis site. We explored the durotactic phenotype of the metastatic MDA-MB-231-Br and -Bo variants on different stiffnesses compared to the parental cells using our previously established fibronectin-coated polyacrylamide (PAA) stiffness gradient gels (stiffness range 0.5-22 kPa)^27^. Briefly, those gradients are made from two PAA solutions resulting in soft and stiff substrates and fluorescent beads are mixed with the stiff solution. One drop of each solution is deposited on a glass-bottom dish and a coverslip is dropped on top, resulting in the mixing of the solutions and the creation of the stiffness gradient at their interface. The measured beads fluorescence intensity from wide-field imaging, correlates with their concentration which increases with the stiffness. While parental cells displayed positive durotaxis and accumulated on the stiff side of the gradients, as shown previously^15^, -Br and -Bo cells were more equally distributed across the gradients from intermediate stiffness to the stiff side (Figure 1A), suggesting a reduced durotaxis range compared to parental cells. Tracking the cells migrating on the soft and stiff side of the gradients confirmed that parental cells preferentially migrate from the soft to the stiff side, while -Br and -Bo cells do not show any direction bias (Extended Data Figures 1A,B,C), indicating an adurotactic phenotype. We then compared -Br and -Bo cell behavior to parental cells on soft (0.5 kPa) and stiff (22 kPa) gels. The metastatic variants spread less than the parental cells on both 22 kPa and 0.5kPa gels (Figures 1C,D). Moreover, on stiff gels, the nuclear translocation of the mechanosensitive transcription factor Yes-associated protein (YAP) was reduced in the -Br and -Bo variants (Figures 1C,D), indicating attenuated mechanotransduction compared to parental cells. In summary, both -Br and -Bo metastatic variants display impaired durotaxis and response to stiffness.

**Figure 1:**
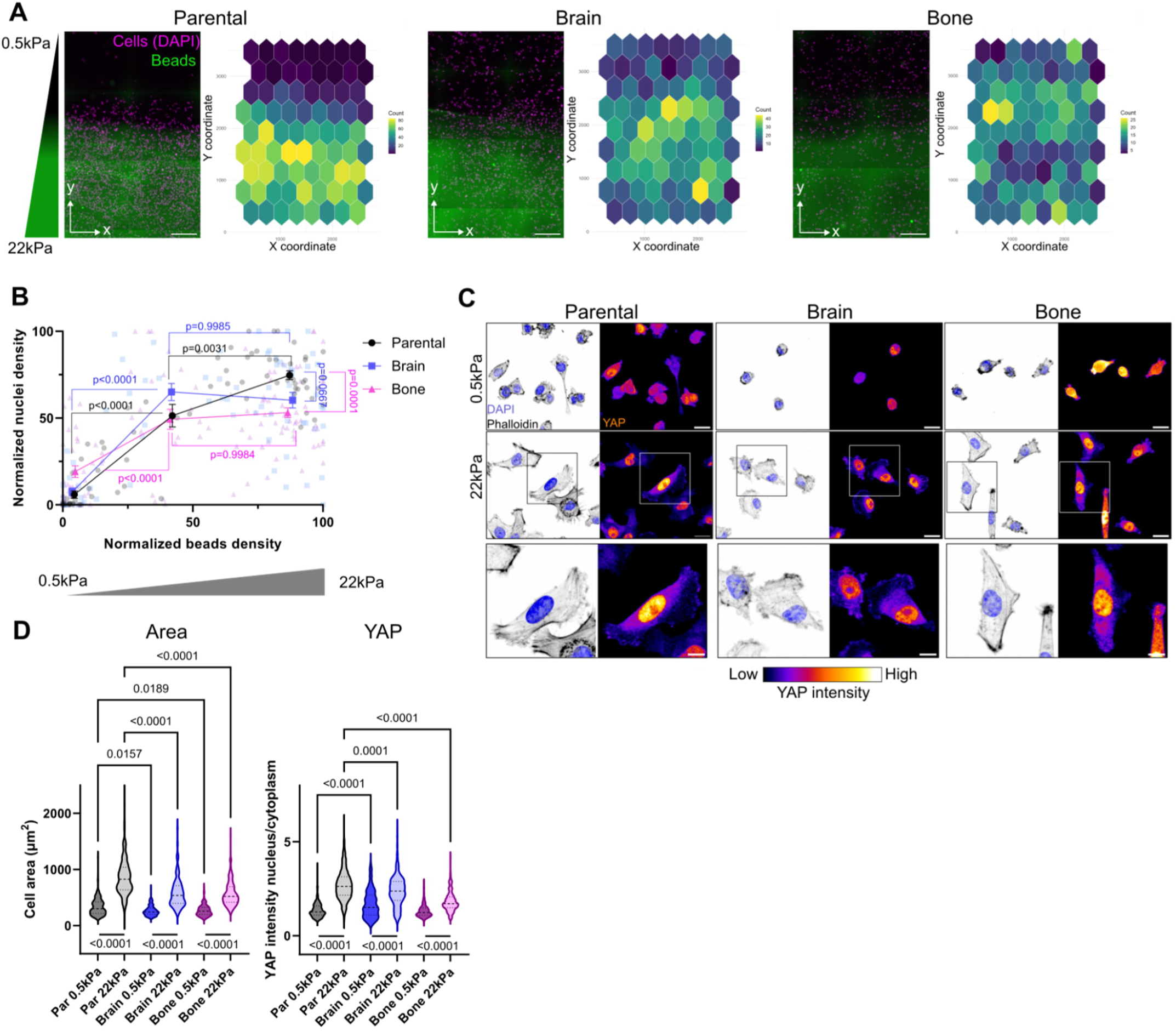
MDA-MB-231 brain and bone metastatic variants have a decreased durotaxis range and impaired response to stiffness. **A)** Representative images of MDA-MB-231 parental, -brain and - bone 3 days after seeding on stiffness gradient gels, the corresponding heat maps showing cell distribution (DAPI, magenta) on the gradients (bead density, green, higher beads density indicates higher stiffness). Scale bar: 500μm. **B)** Quantification of cell densities across the gradients. Data were binned in 3 regions of interest (ROIs) corresponding to soft, intermediate and stiff areas of the gradients. Parental N=72 ROIs from 3 gels, Brain N=72 ROIs from 3 gels, Bone N=120 ROIs from 5 gels, pooled from 3 independent experiments, mean ± standard error of the mean (SEM), ordinary one-way ANOVA with Šídák’s multiple comparisons test. **C)** Representative images of immunofluorescence staining of cells on soft (0.5kPa) and stiff (22kPa) gels (maximum intensity Z projections). Scale bar: 20μm (main), 1μm (insets). **D)** Quantification of cell area and YAP nuclear / cytoplasmic ratio. Median and quartiles are indicated as dashed lines on the violin plots. Cell area: Parental 0.5kPa N=189 cells, 22kPa N=250 cells; Brain 0.5kPa N=131 cells, 22kPa N=249 cells; Bone 0.5kPa N=157 cells, 22kPa N=205 cells. YAP quantification: Parental 0.5kPa N=442 cells, 22kPa N=487 cells; Brain 0.5kPa N=155 cells, 22kPa N=274 cells; Bone 0.5kPa N=200 cells, 22kPa N=463 cells, pooled from 4 independent experiments. Kruskal-Wallis test with Dunn’s multiple comparisons test.

### MDA-MB-231 brain and bone variants form less focal adhesions and have reduced activation of focal adhesion signaling pathways

FAs are the primary cellular mechanosensors, through engagement of the extracellular matrix, and are established regulators of positive durotaxis. We investigated the capacity of -Br and -Bo cells to form FAs and to activate downstream signaling pathways. On stiff gels, both variants formed fewer and slightly smaller FAs than parental cells (Figure 2A and Extended Data Figure 2A). We then assessed the acute activation of signaling pathways downstream of FAs by comparing cells in suspension and cells that have been adhering on fibronectin-coated plastic culture dishes. We observed lower paxillin, focal adhesion kinase (FAK) and AKT phosphorylation levels in adherent -Br and -Bo cells compared to parental cells (Figures 2B,C and Extended Data Figures 2B,C), but no difference in Src and ERK1/2 phosphorylation (Extended Data Figure 2D). To compare the levels of phosphorylated paxillin, FAK and AKT at steady state on different stiffnesses, we analyzed lysates of cells plated for 24h on fibronectin-coated soft (0.5 kPa) and stiff (25 kPa) substrates. We focused on two paxillin phosphorylation sites, Y118 and Y31, both phosphorylated by FAK, enhancing paxillin–FAK interaction and promoting FA turnover^28,29^. On stiff, the Y118 site was less phosphorylated in -Br and -Bo cells compared to parental in two out of four experiments and the Y31 site was consistently less phosphorylated in the metastatic variants compared to parental cells. FAK phosphorylation was reduced on stiff for both variants compared to parental cells, but AKT phosphorylation was reduced on stiff only for -Br cells compared to the parental ones (Figures 2D,E and Extended Data Figure 2E). Taken together, -Br and -Bo cells form fewer FAs and display lower FA signaling to paxillin and FAK in response to cell adhesion and substrate stiffness.

**Figure 2:**
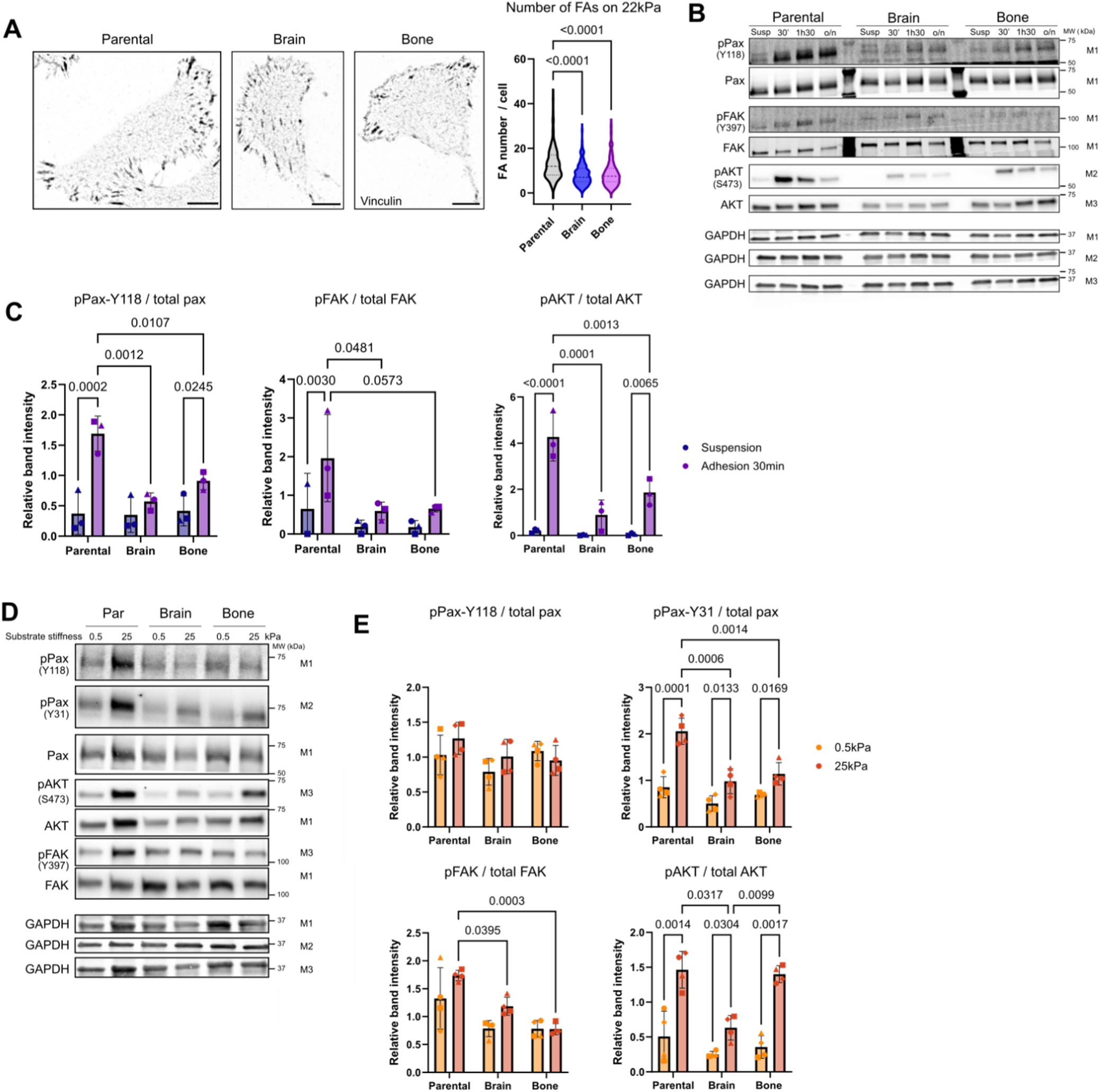
MDA-MB-231 brain and bone variants form less focal adhesions and display lower levels of adhesion-induced paxillin, FAK and AKT phosphorylation. **A)** Representative images of vinculin staining (single plane on the cell surface close to the gel substrate) of cells seeded on 22kPa gels and quantification of the number of focal adhesions (FAs) per cell. Parental N=137 cells, Brain N=143 cells, Bone N=108 cells pooled from 3 independent experiments. Median and quartiles are indicated as dashed lines on the violin plots. Kruskal-Wallis test with Dunn’s multiple comparisons test. Scale bar: 10μm. **B-C)** Representative western blots of the indicated proteins in parental, Br and -Bo MDA-MB-231 cells in suspension (susp) and following adhesion on fibronectin-coated plates (B). GAPDH was used as the loading control. Quantification of the ratio of phosphorylated over total protein is shown for the 30 min timepoint (C). Mean ± standard deviation (SD). N=3 independent experiments. Mixed effects analysis with Tukey’s multiple comparisons test. o/n: overnight. MW: molecular weight. M1-M3: membrane 1-3. **D)** Representative western blots of parental, -Br and -Bo cells after 24h on soft and stiff gels. **E)** Quantification from western blot analyses of the ratio of phosphorylated paxillin, FAK and AKT over the total corresponding proteins comparing cells on soft and stiff gels. N=4 independent experiments. Mean ± SD. 2way ANOVA with Tukey’s multiple comparisons test.

To understand the possible mechanisms underpinning the different durotactic behavior of the parental, -Br and -Bo cells, we employed our comprehensive mass cytometry (MassCytof) antibody panel to gain a profile of the cell-surface receptors, including the primary cell adhesion receptor family, integrins, other established mechanoregulatory adhesion receptors and receptor tyrosine kinases (RTKs)^30^ expressed by these cells. With single-cell resolution, t-distributed stochastic neighbor embedding (t-SNE) analysis revealed that the metastatic variants form overlapping clusters that are distinct from the parental cells (Extended Data Figure 3A), indicating a common cell-surface marker signature between the variants. In particular, -Br and -Bo cells exhibited reduced cell-surface expression of several integrins (Extended Data Figures 3B,C). The lower abundance of integrins on the cell surface is in accordance with the decreased number of FAs observed in these cells (Figure 2A) and may be underpinning their altered mechanosensitive behavior.

### Soft-conditioned MDA-MB-231 cells respond similarly than brain and bone variants to stiffness increase

During in vivo selection, the -Br and -Bo cells were enriched not only for their ability to grow in environments of different stiffness, but also for their ability to survive in the blood circulation and to exit the blood vessels to home into specific tissues. To identify the phenotypes and mechanisms specific for their response to stiffness, we cultured MDA-MB-231 on soft fibronectin-coated gels of 0.5kPa for six or more passages and compared their behavior on stiffness gradients and on soft and stiff gels with parental cells. On 0.5 to 22kPa gradients, the soft-conditioned cells were equally distributed from intermediate stiffness to stiff, thus resembling the behavior of the metastatic variants (Figures 3A,B). Moreover, the soft-conditioned cells were less spread and had less nuclear YAP compared to parental cells on stiff gels (Figures 3C,D). They also formed less FAs on stiff (Figure 3E), and had lower levels of paxillin and FAK phosphorylation on stiff but showed no difference in AKT activation (Figures 3F,G and Extended Data Figure 4). Taken together, these data indicate that the soft-conditioned parental MDA-MB-231 cells phenocopy the dampened mechanoresponses observed in the -Br and -Bo variants.

**Figure 3:**
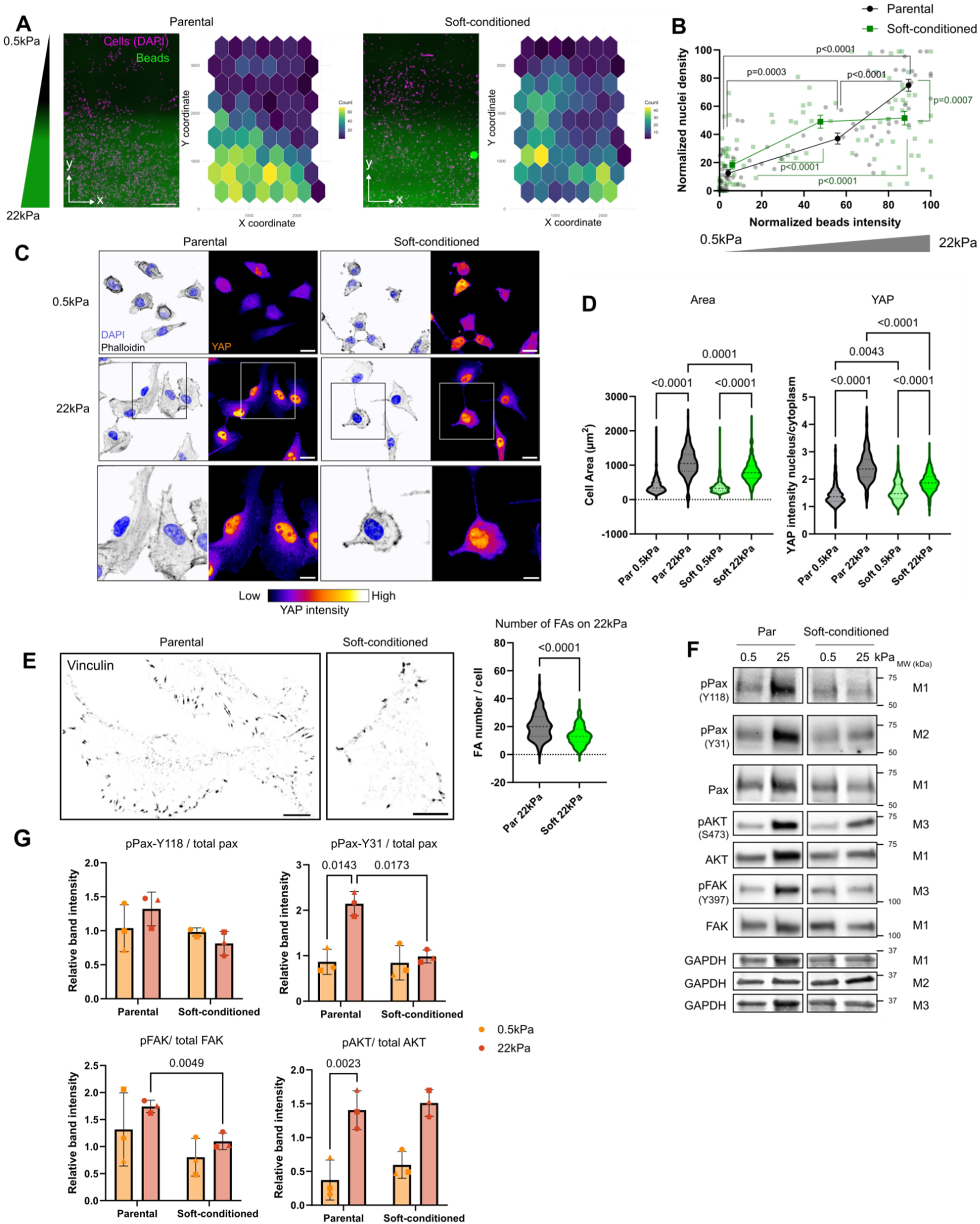
Soft-conditioned MDA-MB-231 cells show similar durotaxis, stiffness response and focal adhesion signaling to the MDA-MB-231 brain and bone variants. **A)** Representative images of MDA-MB-231 parental and soft-conditioned cells 3 days after seeding on stiffness gradient gels and corresponding heat maps showing their distribution on the gradients. Scale bar: 500μm. **B)** Quantification of cell densities across the gradients. Data were binned in 3 regions of interest corresponding to soft, intermediate and stiff areas of the gradients. Parental N=72 ROIs from 3 gels, soft-conditioned N=96 ROIs from 4 gels, pooled from 2 independent experiments, mean ± SEM, ordinary one-way ANOVA with Šídák’s multiple comparisons test. **C)** Representative images of immunofluorescence staining of cells on soft and stiff gels (maximum intensity Z projections). Scale bar: 20μm. **D)** Quantification of cell area and YAP nuclear / cytoplasmic ratio. Median and quartiles are indicated as dashed lines on the violin plots. Cell area: parental 0.5kPa N=381 cells, 22kPa N=232 cells; soft-conditioned 0.5kPa N=308 cells, 22kPa N=244 cells from 5 independent experiments. YAP quantification: parental 0.5kPa N=362 cells, 22kPa N=262 cells; soft-conditioned 0.5kPa N=267 cells, 22kPa N=255 cells pooled from 4 independent experiments. Kruskal-Wallis test with Dunn’s multiple comparisons test. **E)** Representative images of vinculin staining (single plane on the cell surface close to the gel substrate) of cells seeded on 22kPa gels and quantification of the number of focal adhesions per cell. Parental N=176; soft-conditioned N=186 cells pooled from 5 independent experiments. Median and quartiles are indicated as dashed lines on the violin plots. Mann-Whitney test, two-tailed. **F)** Representative western blots of parental and soft-conditioned cells after 24h on soft and stiff gels. MW: molecular weight. M1-M3: membrane 1-3. **G)** Quantification from western blot analyses of the ratio of phosphorylated paxillin, FAK and AKT over the total corresponding proteins comparing cells on soft and stiff gels. N=4 independent experiments. Mean ± SD. Two-way ANOVA with Uncorrected Fisher’s LSD.

### Leupaxin is more expressed in MDA-MB-231-Br and -Bo compared to parental and localizes to paxillin-low FA subdomains

To further investigate, in an unbiased way, which genes are differentially expressed in the metastatic variants compared to parental cells, we performed single-cell RNA sequencing. Louvain clustering indicated that the three cell lines formed separate clusters without clear overlap (Figure 4A). In addition, we computed diffusion maps and pseudotime for the parental-metastatic trajectories. Weak connectivity between populations in the diffusion maps further supported distinct transcriptomic features (Figure 4B). This suggests that, in contrast to arising from the original population, metastatic variant selection has involved transcriptomic reprogramming of parental cells to enable adaptation to stressors such as barriers in the primary tumor and blood circulation, and establishment of metastatic lesions in the secondary organs. Upon studying the parental-metastasis trajectories the features most closely associated with a transcriptomic shift towards the metastatic variants were upregulation of *LAMB3*, *G0S2*, *PLAUR, SH3BGRL, EMP1, LPXN* and *MMP14* (Figure 4C). Of these, *MMP14* and *SH3BGRL* have been previously linked to metastasis and other pro-oncogenic properties have been described for *G0S2*, *PLAUR, EMP1 and LPXN* in breast cancer^31–36^. We chose to focus on leupaxin (LPXN), a LIM-domain protein belonging to the paxillin family^37,38^ as its role in mechanosensing has not been studied despite its localization to FAs^39^. Furthermore, we found a correlation between high leupaxin expression and a previously published lymphovascular invasion signature (Extended Data Figure 5A)^40^, implying a possible role for it in the context of metastasis. Western blot analysis confirmed higher leupaxin protein expression in both the -Br and -Bo variants. Other FA regulatory proteins such as talin-1, kindlin-1 and -2 and paxillin were either unchanged or only modestly affected in one of the variants (kindlin-1 in -Br) (Extended Data Figure 5B). Moreover, we found LPXN to be also more expressed in the soft-conditioned cells compared to parental cells, with the same level of expression as the -Bo cells, both on soft and stiff (Figures 4D,E). Next, we explored leupaxin and paxillin FA localization in parental, -Br and -Bo cells. Enhanced super-resolution radial fluctuations (eSRRF) image reconstruction, which exploits time series images with fluctuating fluorescence signal (here from spinning disk confocal imaging) was used to reconstruct super-resolution images^41^. The imaging showed higher leupaxin staining, relative to paxillin, in the FAs of the variants. Furthermore, leupaxin and paxillin showed alternating localization within FAs with leupaxin occupying FA subdomains with low paxillin signal (Figure 4E). This almost mutually exclusive patterning between leupaxin and paxillin suggested that leupaxin could be competing with paxillin and, thus, impact FA formation and signaling and durotaxis in the -Br, -Bo and soft-conditioned cells where it is highly expressed.

**Figure 4:**
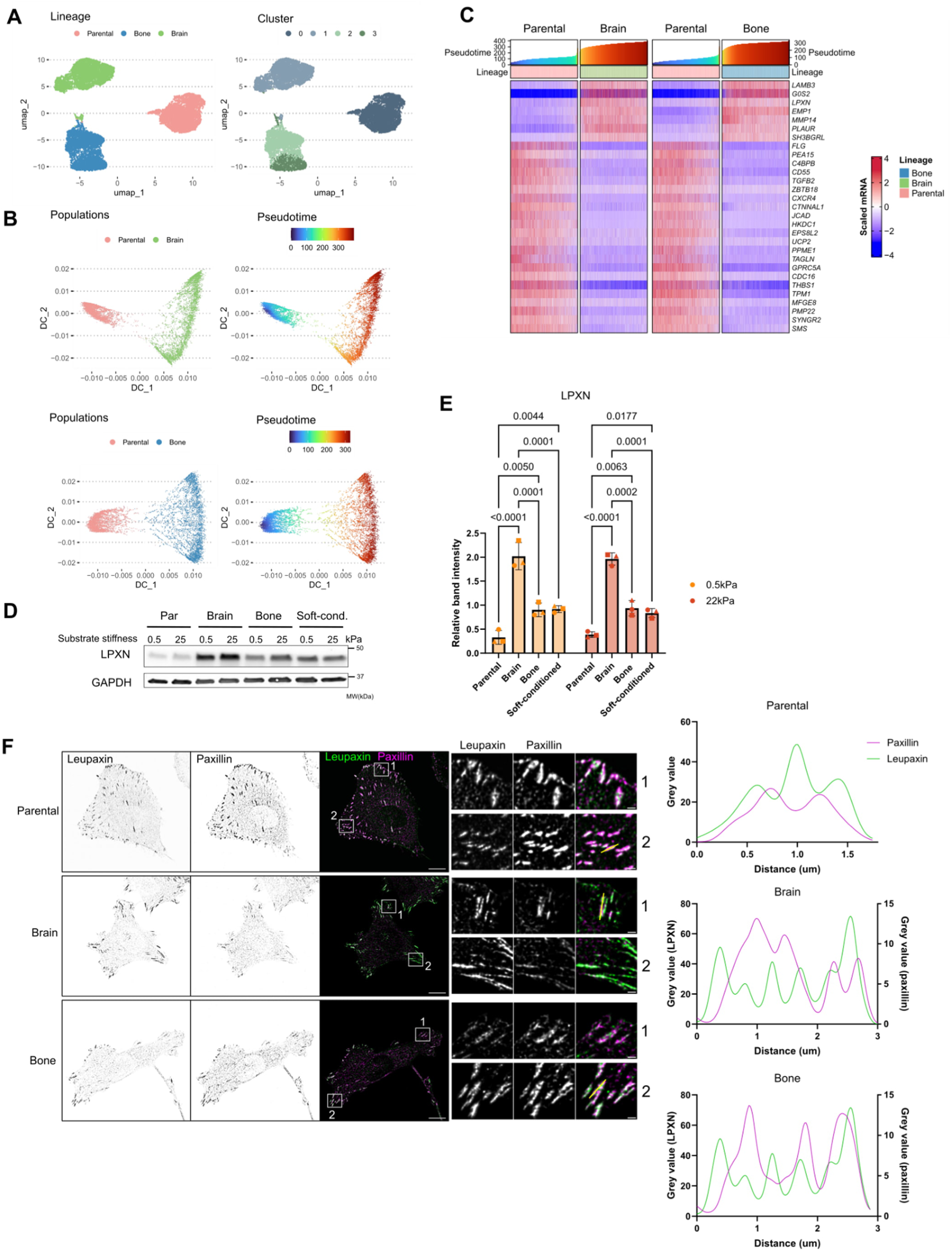
MDA-MB-231 brain and bone variants share transcriptomic similarities with soft-conditioned MDA-MB-231 and express high levels of leupaxin. **A)** Louvain clustering and UMAP visualization of single-cell RNA sequencing of parental, -Br and -Bo cells. The left graph shows the different cell lines, the right graph shows the Louvain clusters. **B)** Diffusion map and pseudotime analysis of the single-cell RNA sequencing data. The different clusters are shown on the left graph and the pseudotime scale is indicated on the right graph. **C)** Most robustly correlating shared features of the parental-metastasis trajectories in -Br and -Bo cells. **D**-**E)** Representative western blot of parental, -Br, -Bo and soft-conditioned cells showing LPXN expression on soft and stiff gels (D), and corresponding quantification (E). Mean ± SD. N=3 independent experiments. Two-way ANOVA with Šídák’s multiple comparisons test. **F)** Representative immunofluorescence images of leupaxin and paxillin staining in parental, -Br and -Bo cells seeded on fibronectin-coated glass-bottom dishes. Image resolution was enhanced by eSRRF processing. Graphs on the right indicate the paxillin and leupaxin intensity along the yellow lines indicated on the insets. Scale bars: 10μm (main), 1μm (inset).

### Highly expressed leupaxin impairs durotaxis and FA formation

To test if high leupaxin expression is implicated in the observed stiffness response phenotypes, we silenced leupaxin in -Br cells. On stiffness-gradient gels, LPXN-silenced (siLPXN) -Br cells reverted to the parental cell behavior, accumulating on the stiff side of the gels (Figure 5A), indicating that leupaxin is responsible for the impaired durotaxis of the -Br cells. In parallel, western blot analysis of cell lysates harvested from the same experiments showed an increase in paxillin phosphorylation in siLPXN cells, but no difference in FAK phosphorylation (Extended Data Figure 6A), indicating that leupaxin inhibits paxillin activation. To test whether leupaxin alone is sufficient to switch the durotaxis and signaling behavior of the cells, we generated MDA-MB-231 cells stably overexpressing GFP-leupaxin (GFP-LPXN), sorted based on high and low GFP-LPXN expression (referred to as GFP-LPXN high and GFP-LPXN low) (Extended Data Figure 6B). We confirmed in the GFP-LPXN high cells that the GFP-LPXN localizes to FAs (Extended Data Figure 6C) and had a similar mutually exclusive patterning with paxillin as observed with the endogenous protein. On stiffness-gradient gels, the GFP-LPXN high cells were similar to the -Br and -Bo cells, being equally distributed from the intermediate to the stiff side of the gradient (Figure 5B). The GFP-LPXN effect was concentration dependent as the GFP-LPXN low cells were not significantly different from the parental cells (Extended Data Figure 6D), in line with the notion of leupaxin competing with paxillin to inhibit mechanosensing. We did not observe any difference in cell spreading and YAP nuclear localization on stiff gels (Figures 5C,D). However, on soft, the GFP-LPXN high cells were slightly larger and had more nuclear YAP than the parental cells (Figures 5C,D), suggesting that high LPXN expression enables cells to adapt better to softer surfaces. Moreover, the GFP-LPXN high cells formed less FAs on stiff than the parental cells (Figure 5E), indicating that leupaxin overexpression is sufficient to induce a decreased durotaxis range, to inhibit FA formation on stiff and enhance nuclear YAP translocation in cells on soft. Taken together, these data confirm the role of leupaxin as a negative regulator of durotaxis and a mechanomodulator supporting adhesion-dependent processes on soft.

**Figure 5:**
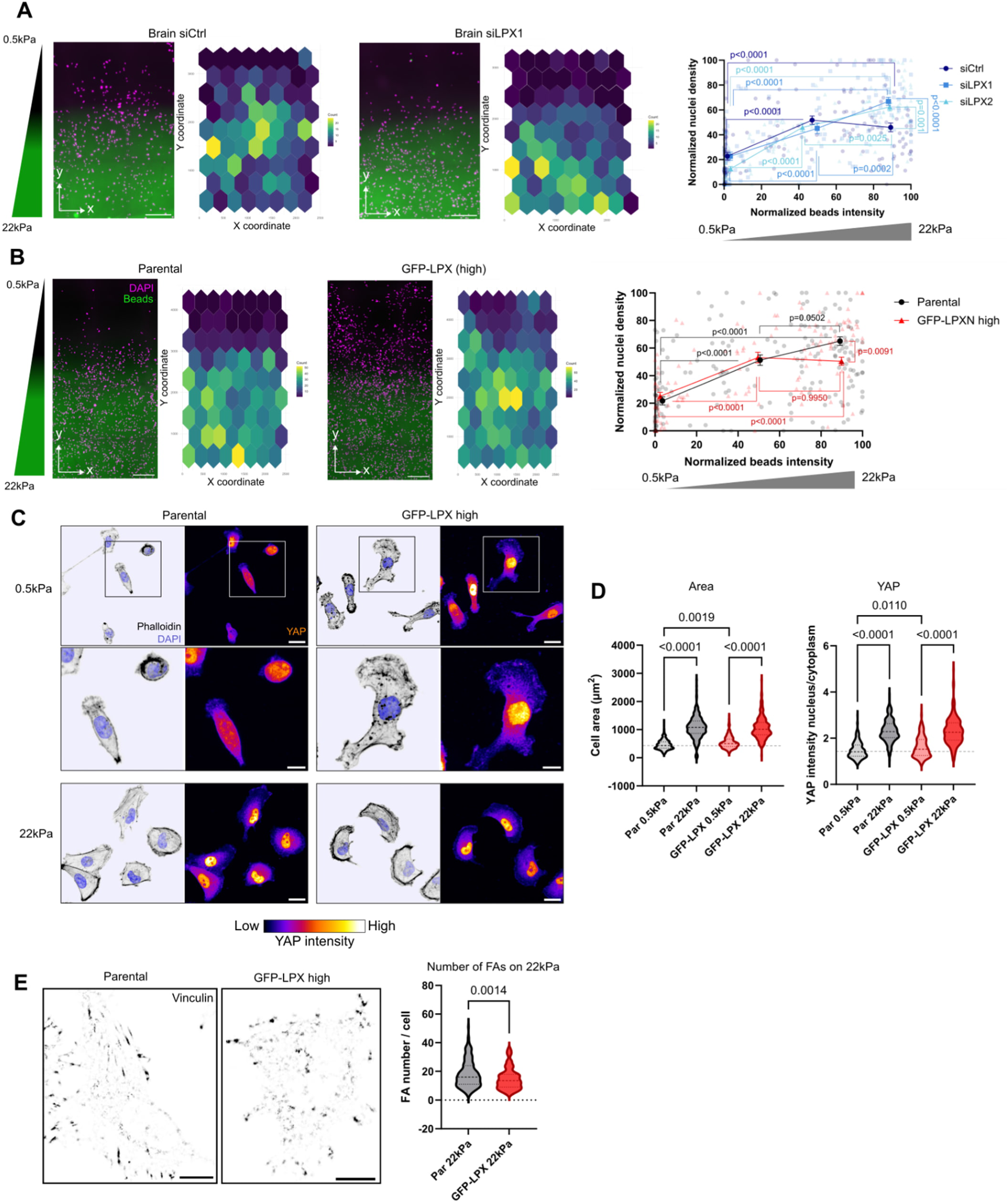
Manipulating leupaxin levels impacts durotaxis and mechanosensing. **A)** Representative images of MDA-MB-231-Br transfected with non-targeting siRNA (siCtrl) and LPXN siRNA-1 three days after seeding on stiffness gradient gels, the corresponding heat maps showing their distribution on the gradients and quantification of cell densities across the gradients for siCtl, siLPXN1 and siLPXN2 conditions. Data were binned in 3 regions of interest corresponding to soft, intermediate and stiff areas of the gradients. siCtrl N=144 ROIs from 6 gels, siLPXN1 N=120 ROIs from 5 gels, siLPXN2 N=144 ROIs from 6 gels, pooled from 3 independent experiments, mean ± SEM, ordinary one-way ANOVA with Šídák’s multiple comparisons test. Scale bar: 500μm. **B)** Representative images of MDA-MB-231-parental and stably expressing GFP-LPXN at high level of expression 3 days after seeding on stiffness gradient gels, corresponding heat maps showing their distribution on the gradients and quantification of cell densities across the gradients. Data were binned in 3 regions of interest corresponding to soft, intermediate and stiff areas of the gradients. Parental N=136 ROIs from 5 gels, GFP-LPXN high N=184 ROIs from 7gels pooled from 5 independent experiments, mean ± SEM, ordinary one-way ANOVA with Šídák’s multiple comparisons test. Scale bar: 500μm. **C)** Representative images of immunofluorescence staining of parental and GFP-LPXN high cells on soft and stiff gels (maximum intensity Z projections). Scale bar: 20μm. **D)** Quantification of cell area and YAP nuclear / cytoplasmic ratio. Median and quartiles are indicated as dashed lines on the violin plots. Cell area: parental 0.5kPa N=276 cells, Parental 22kPa N=220 cells, GFP-LPXN 0.5kPa N=290 cells, GFP-LPXN 22kPa N=205 cells from 4 independent experiments. YAP quantification: parental 0.5kPa N=252 cells, Parental 22kPa N=200 cells, GFP-LPXN 0.5kPa N=251 cells, GFP-LPXN 22kPa N=196 cells pooled from 3 independent experiments. Kruskal-Wallis test with Dunn’s multiple comparisons test. **E)** Representative images of vinculin staining (single plane on the cell surface close to the gel substrate) of cells seeded on 22kPa gels and quantification of the number of focal adhesions per cell. Parental 22kPa N=166 cells, GFP-LPXN 22kPa N=144 cells pooled from 4 independent experiments. Median and quartiles are indicated as dashed lines on the violin plots. Mann-Whitney test, two-tailed. Scale bar: 10μm.

### Leupaxin impairs durotaxis by inhibiting the paxillin-FAK pathway

Next, we investigated if leupaxin alters cell signaling as a function of stiffness. GFP-LPXN high cells, plated on 0.5 and 25 kPa substrates, had lower levels of phosphorylated paxillin and FAK, but higher levels of phosphorylated AKT compared to the parental and GFP-LPXN low cells (Figures 6A,B and Extended Data Figure 7A). This suggested that leupaxin might impair durotaxis via inhibiting the paxillin–FAK signaling axis. To test this, we treated parental cells on stiffness gradients with the specific FAK inhibitor ifebemtinib^42,43^. Ifebemtinib efficiently inhibited FAK phosphorylation (Extended Data Figure 7B) and significantly inhibited positive durotaxis at the higher stiffness range resulting in similar distribution of cells across the gradient, phenocopying the behavior of GFP-LPXN high cells and the metastatic variants (Figures 6C,D). Moreover, as active paxillin interaction with FAK has recently been shown to drive positive durotaxis^8^, we performed proximity ligation assays (PLA) between phosphorylated paxillin-Y31 and FAK in parental and GFP-LPXN high cells. Leupaxin overexpression resulted in a decrease of PLA dots density (Figure 6E and Extended Data Figure 7D), indicative of decreased interaction between active paxillin and FAK. Taken together, these data indicate leupaxin as the first negative regulator of durotaxis, which acts by modulating the paxillin–FAK pathway.

**Figure 6:**
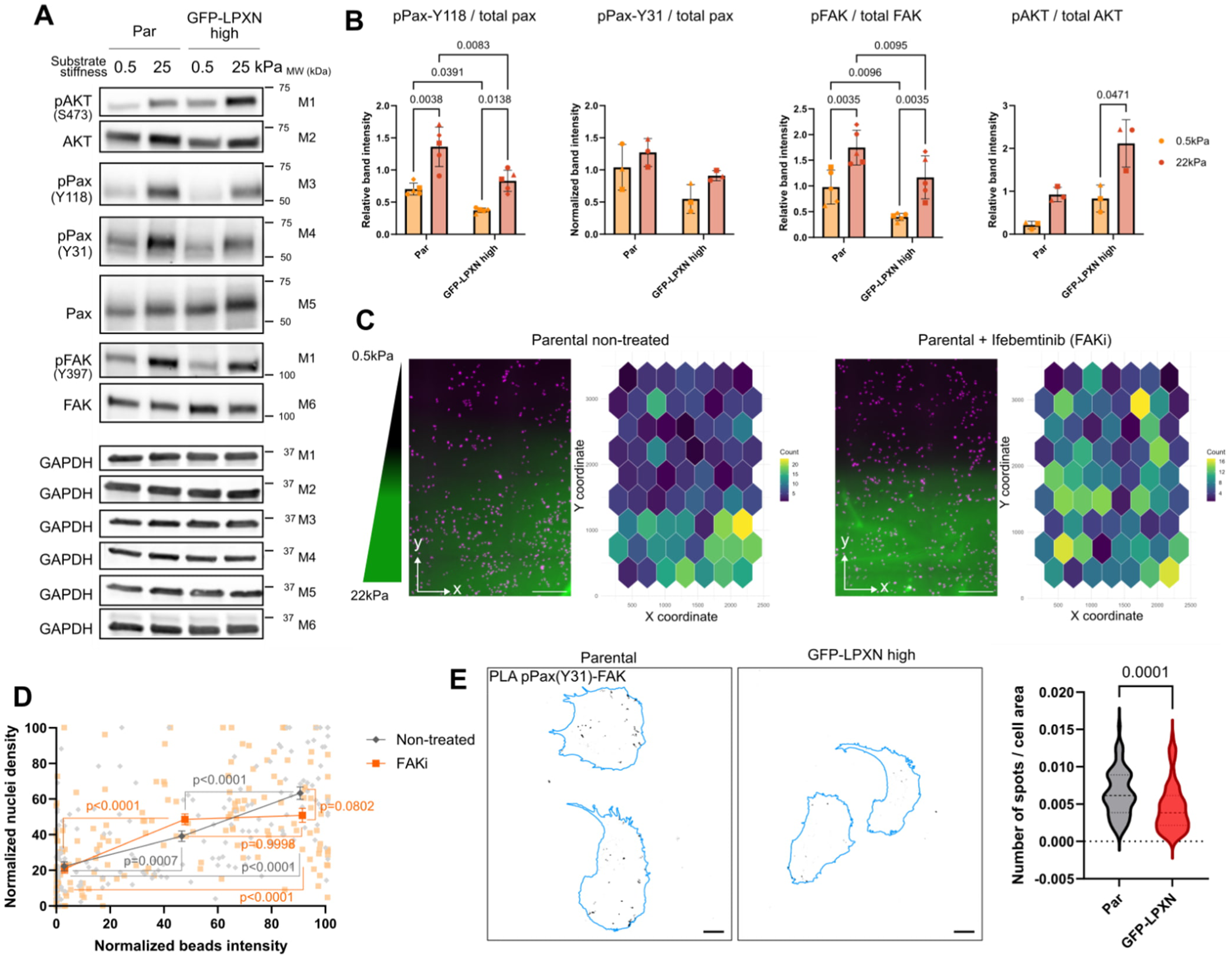
Leupaxin modulates durotaxis by inhibiting paxillin and FAK. **A)** Representative western blots of parental and GFP-LPXN high and low expressing cells after 24h on soft and stiff gels. **B)** Quantification from western blot analyses of the ratio of phosphorylated paxillin, FAK and AKT over the total corresponding proteins comparing cells on soft and stiff gels. pPaxillin-Y118 and pFAK: N=5 independent experiments, two-way ANOVA with Uncorrected Fisher’s LSD. Mean ± SD. **C)** Representative images of MDA-MB-231 parental untreated and treated with 0.001μM Ifebemtinib 3 days after seeding on stiffness gradient gels and corresponding heat maps showing their distribution on the gradients. Scale bar: 500μm. **D)** Quantification of cell densities across the gradients. Data were binned in 3 regions of interest corresponding to soft, intermediate and stiff areas of the gradients. N=168 ROIs from 7 gels pooled from 3 independent experiments, mean ± SEM, ordinary one-way ANOVA with Šídák’s multiple comparisons test. **E)** Representative images of pPax-Y31 and FAK PLA in MDA-MB-231 parental and GFP-LPXN high expression. The cell outlines, drawn from phalloidin labelling, and shown in blue. The graph on the right side shows the quantification of the number of spots in each cell divided by the corresponding cell area. Parental: N=77 cells, GFP-LPXN: N=90 cells pooled from 3 independent experiments. Median and quartiles are indicated as dashed lines on the violin plots. Unpaired t-test, two-tailed. Scale bar 20μm.

## Discussion

In this study, we unveil a new mechanism regulating durotaxis in metastatic breast cancer cells. We identify elevated leupaxin expression as a shared feature between the MDA-MB- metastatic variants and cells conditioned by long-term exposure to a soft culture environment. Through loss and gain-of-function experiments we demonstrate that leupaxin is necessary and sufficient to attenuate positive durotaxis at higher stiffness regimes and that leupaxin enables metastatic cells to adapt to a wider stiffness range.

Unlike paxillin, the role of leupaxin in adhesion regulation remains incompletely understood. It has been previously shown to localize to FAs in MDA-MB-231 cells^39^. In HEK293T cells, leupaxin FA localization has been shown to be dependent on its LIM3 domain and to inhibit paxillin phosphorylation^44^. Leupaxin-mediated inhibition of paxillin phosphorylation was also shown in podosomes of RAW 246.7 cells (a mouse macrophage/monocyte-like cell line)^45^. However, its effect on downstream FA signaling and its role in mechanosensing and durotaxis had not been investigated before. We showed that leupaxin localizes in paxillin-low FA subdomains in MDA-MB-231 cells and inhibits paxillin and FAK phosphorylation when highly expressed as in the -Br and - Bo metastatic variants. Our data are supportive of leupaxin competing with paxillin and reducing transmission of mechanical information through signaling. Durotaxis and mechanosensing are likely to be regulated by the balance of different FA components which determine FA stability, such as the leupaxin-paxillin ratio. Moreover, FAK is a well know actor of mechanotransduction and positive durotaxis^2,5,46–48^. We find that leupaxin inhibits integrin-mediated FAK activation and durotaxis, indicating that leupaxin is a key inhibitor of the paxillin-FAK axis underpinning its ability to disrupt durotaxis. Interestingly, leupaxin is highly expressed in immune cells such as B-cells and macrophages^37,49^, which characteristically do not generate strong FAs or stress fibers and were recently demonstrated to show no durotaxis^8^. This implies that leupaxin may be an important durotaxis inhibitor across various cell types. The cell- and tissue-type-specific expression patterns of leupaxin and its contribution to durotaxis and organotropic metastasis to soft tissue are important topics for future research.

The mechanisms regulating positive and negative durotaxis remain to be fully understood. Positive durotaxis mechanisms have been more widely studied^50^ with integrin adhesion complexes (IACs), FAK^2,46,47,51,52^ and the actin cytoskeleton^5,53,54^ identified as the main actors. In contrast, while modulating FA reinforcement^15,55^ and cell contractility^15,17^ can switch cell migration between positive and negative durotaxis or adurotaxis, the genes regulating which phenotype is predominant have not been identified. Therefore, our discovery of leupaxin as a key determinant of durotaxis phenotypes is a significant discovery.

According to the molecular clutch model, which involves myosin motors pulling actin fibers connected to substrate engaging IACs (clutches), cells migrate toward their stiffness optimum^56–58^. As most cell types, grown in vitro, display monotonously increasing forces with increasing stiffness, they undergo positive durotaxis. This has been attributed to vinculin and talin-dependent FA reinforcement. However, when FA reinforcement is lacking, a biphasic force regime is observed where cells mount maximum forces at intermediate stiffnesses^55^ and, as shown by us, display either negative or positive durotaxis and migrate up or down a gradient towards their stiffness optimum^15^. FA dynamics and FAK activity are emerging as key determinants of cells durotaxis phenotype. FAK activity is linked to FA turnover^29^. Signaling via the paxillin-FAK axis drives positive durotaxis in pancreatic cancer^8^ and we observed that FAK inhibition by leupaxin or FAK kinase inhibitor leads to attenuated positive durotaxis. A better understanding of how fine tuning of FA dynamics by various regulators beyond FAK affect durotaxis could be of interest to understand durotaxis regulation.

Adurotaxis has been previously observed for a subset of MDA-MB-231 cells^17^. Our findings that variants of MDA-MB-231 have an adurotactic behavior and decreased sensitivity to stiffness are concordant with this earlier observation. Yeoman et al., showed that weakly adherent cells are more adurotactic than strongly adherent ones. This is also in line with our finding that leupaxin, as a negative regulator of FAs, drives this phenotype. Moreover, Yeoman et al. showed that increased contractility in weakly adherent cells explains their adurotaxis. We did not explore cell contractility and actin cytoskeleton, but further investigation on the interplay between leupaxin, cell contractility and durotaxis would be of interest.

Decreased positive durotaxis and mechanosensing could contribute to tumor cell invasion and their ability to proliferate in an environment with different mechanical properties than the primary tumor. Indeed, it was recently shown that in mice, breast tumor xenografts include soft-niches from which brain metastatic cells emerge^59^. Similarly, we found similarities between soft-conditioned cells and the metastatic variants. We believe that the loss of mechanosensitivity in these cells induces the observed impaired durotaxis. Moreover, the similarities observed between the soft-selected cells and the metastatic variants indicate that cell plasticity and adaptation to alternating stiffness and adhesion landscapes are essential during the metastatic cascade. This notion is in line with the previous observed similarities between the -Br cells and cells from soft niches of breast tumors xenografts^59^, which suggests that metastatic cells emerging from the primary tumor could share features of defective durotaxis.

Interestingly, leupaxin has been identified as an oncogene and was shown to promote invasion in prostate cancer^60^, bladder cancer^61^ and breast cancer^36^. This, and the correlation we found between high leupaxin expression and lymphatic vessel invasion signature in invasive breast cancer, imply that leupaxin-driven defects in cell mechanoresponse could be an important factor of metastasis formation in triple-negative breast cancer. Further studies will be needed to understand the clinical relevance of leupaxin-driven durotaxis alteration.

## Material and methods

### Cell lines and cell culture

MDA-MB-231-Brain and -Bone metastatic variants where primarily generated from MDA-MB-231 parental cells and obtained from Dr. Joan Massagué’s^23^ and Dr. Riko Nishimura’s^22^ laboratories respectively. They were authenticated as MDA-MB-231 using a short tandem repeat assay (Leibniz Institute DSMZ—German Collection of Microorganisms and Cell Cultures). Cells were regularly tested for mycoplasma contamination and were cultured in Dulbecco’s Modified Eagles Medium (DMEM) high glucose (VWR 392-0413) with 10% fetal bovine serum, 1% L-glutamine (Sigma-Aldrich), 1% penicillin-streptomycin (Sigma-Aldrich) and 1% MEM non-essential amino acids (Sigma-Aldrich) at 37°C in 5% CO_2_ atmosphere.

For cell culture on soft dishes or 6-well plates, 0.5kPa PetriSoft Easy Coat 10cm dishes or 0.5 and 25kPa SoftWell Easy Coat 6-well plates (Cell Guidance Systems) where coated overnight with fibronectin (bovine plasma, Sigma-Aldrich) at 10µg/mL in PBS overnight at 4°C before seeding cells.

Soft-conditioned cells were generated by seeding MDA-MB-231 parental cells on 0.5kPa fibronectin-coated dishes and were used for experiments after more than 6 passages on those dishes.

For GFP-LPXN stable overexpression, the human leupaxin sequence fused to GFP in N-terminal by a linker (SLGGSRA) (gBlock gene fragment from Integrated DNA Technologies) was cloned in a pPB-CAG piggyback vector. MDA-MB-231 parental cells were transfected with the pPB-GFP-LPXN plasmid using Lipofectamine 3000 (LifeTechnologies). Successfully transfected GFP positive cells where isolated by FACS based on GFP levels of expression to generate high and low overexpressing cells.

For LPXN silencing, cells at 80% confluence in a 6-well plate containing 2mL of growth medium without antibiotics were transfected using 9uL of Lipofectamine RNAiMax (Invitrogen) with 30pmol of ON-TARGETplus Non-targeting siRNA #1 (Horizon Discovery D-001810-01-05) or 15pmol of ON-TARGETplus Human LPXN siRNA (Horizon Discovery J-009746-06-0002: siLPXN1, sequence CUUCGGAGAUCCUUUCUAU and J-009746-07-0002: siLPNX2, sequence GCGCAGCUCGUGUAUACUA) diluted in 300uL OptiMEM (Gibco 31985054). Media was changed 24h after transfection and cells were seeded on stiffness gradient gels 48h after transfection, while remaining cells were seeded on 6-well plates for cell lysis at the end point of the experiment 3 days later.

### Polyacrylamide hydrogels

Polyacrylamide gels of different stiffnesses, either homogenous or with a stiffness gradient, were generated as previously described^27^. 35mm #1 glass-bottom dishes (Cellvis) were treated for 30min with 100µL Bind-Silane solution (3-(trimethoxysilyl)propylmethacrylate (7.15% by volume, Sigma-Aldrich, M6514) and acetic acid (7.15% by volume) in absolute ethanol). The dishes were then washed twice with absolute ethanol and left to dry. 40% (w/v) acrylamide and 2% (w/v) N,N-methyl-bis-acrylamide where mixed in PBS at appropriate ratios to obtain the desired stiffness (Table 1) and kept on ice. For stiffness gradient gels, the solution corresponding to 22kPa was supplemented with 0.1 µm yellow-green or red fluorescent (505/515 or 580/605) microspheres (∼1.2 × 10^11^/mL final concentration, Invitrogen, F8803 or F8801) that were sonicated for 3min. 10% ammonium persulfate (final 0.1% by volume, Bio-Rad) and N,N,N′,N′-tetramethylethylenediamine (final 0.2% by volume, Sigma-Aldrich, T-9281) were then added to initialize the polymerization. For homogenous gels, a 15.6 µL drop was immediately deposited on the glass and covered by a 13mm coverslip. For stiffness-gradient gels, 7.8 µL drops of each solution (0.5 and 22kPa) were deposited and left to diffuse together under the coverslip. After 1h15, PBS was added on top and the coverslips were carefully removed.

**Table 1:**
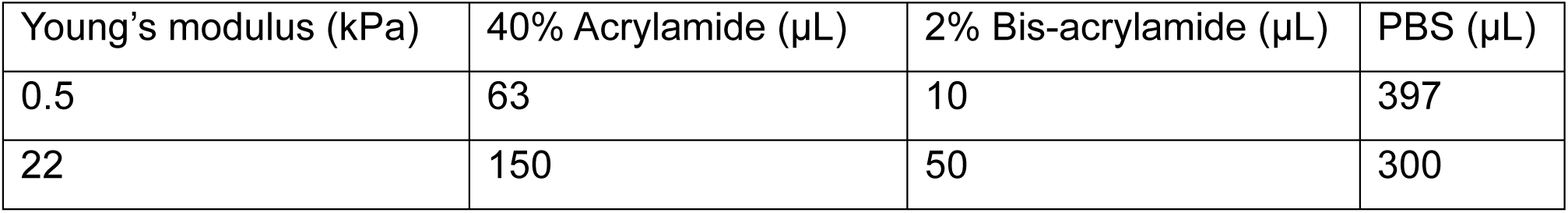
PAA mixes to prepare homogenous and stiffness gradient gels.

Before use, the hydrogels were activated by 500µL of a solution of 0.2 mg ml–1 Sulfo-SANPAH (Thermo Fisher Scientific, 22589) and 2 mg ml–1 N-(3-dimethylaminopropyl)-N′ ethylcarbodiimidehydrochloride (Sigma-Aldrich, 03450) in 50 mM 4-(2 hydroxyethyl)piperazine-1-ethanesulfonic acid. The gels were left in gentle agitation protected from light at room temperature during 30min before irradiation with UV (28–32 mW/cm^2^) for 10min. The dishes were then washed three time with PBS and coated with 10µg/mL fibronectin in PBS overnight at 4°C.

### Migration experiments on stiffness gradient gels

20 000 cells were seeded in each dish containing the fibronectin-coated gels. One hour later, even seeding through the gradient was controlled visually by bright-field microscopy. The cells were then either used for live imaging after 24h or fixed with 4% paraformaldehyde (PFA) after 72h and stained with 5 µg/mL DAPI.

### Immunofluorescence staining

20 000 cells were seeded on homogenous 0.5 or 22kPa PAA gels and fixed 72 hours later. The cells were permeabilized in 0.1% Triton-X100 in PBS for 5min and blocked with 10% horse serum in PBS for 30min. Primary antibodies were incubated in PBS 10% horse serum overnight at 4°C. Secondary antibodies were incubated in PBS 10% horse serum for 1h at room temperature. 0.033nM Phalloidin-atto647N (Sigma-Aldrich) was added together with the secondary antibodies. The nuclei were stained with 5 µg/mL DAPI for 15min.

### Proximity ligation assay (PLA)

30 000 cells were seeded on fibronectin-coated glass-bottom dishes (Cellvis), fixed 24h later in 4% PFA and 0.3% Triton-X100 in PBS for 10min and blocked in 5% horse serum and 0.3% Triton-X100 for 15min. Primary antibodies (see Table 2) in PBS and 5% horse serum were then incubated with the cells at room temperature for 1h15 and washed with PBS. Duolink In Situ PLA Probe Anti-Mouse PLUS (Sigma-Aldrich DUO92001) and Duolink In Situ PLA Probe Anti-Rabbit MINUS (Sigma-Aldrich DUO92005) were then incubated at room temperature for 50min, followed by a 30min incubation at 37°C and two 5min washes in wash buffer A (Sigma-Aldrich DUO82049), immerging fully the dishes in wash buffer under agitation. Cells were then incubated with ligase in ligation buffer (from Duolink In Situ Detection Reagents Red, Sigma-Aldrich DUO92008) for 30min at 37°C and washed twice with wash buffer A. The amplification mix containing the polymerase in amplification buffer (from Duolink In Situ Detection Reagents Red, Sigma-Aldrich DUO92008) was then incubated for 100min at 37°C. The dishes were washed twice for 10min with wash buffer B (Sigma-Aldrich DUO82049) and with wash buffer A for 1min. 0.033nM Phalloidin-atto647N (Sigma-Aldrich) and 5 µg/mL DAPI were then incubated for 45min and washes twice for 2min with wash buffer A and once for 1min with wash buffer B, and finally kept in PBS for imaging.

**Table 2:** antibodies for immunofluorescence, Western blot and PLA.

| Antibody | Company | Reference | Dilution used |
| --- | --- | --- | --- |
| **Immunofluorescence** |  |  |  |
| Anti-YAP1 antibody [EP1674Y] Rabbit | Abcam | ab52771 | 1:100 |
| Anti-Vinculin Antibody mouse monoclonal, hVIN-1 | Sigma-Aldrich | V9131 | 1:500 |
| Donkey anti-Rabbit IgG (H+L) Highly Cross-Adsorbed Secondary Antibody, Alexa Fluor™ 488 | Invitrogen | A-21206 | 1:300 |
| Donkey anti-Mouse IgG (H+L) Highly Cross-Adsorbed Secondary Antibody, Alexa Fluor™ 568 | Invitrogen | A10037 | 1:300 |
| **PLA** |  |  |  |
| Phospho-Paxillin (Tyr31) Polyclonal Antibody Rabbit | Invitrogen | 44-720G | 1:250 |
| Purified Mouse Anti-FAK Clone 77/FAK | BD Transduction Laboratories | 610088 | 1:100 |
| **Western blot** |  |  |  |
| Phospho-Paxillin (Tyr118) Antibody, rabbit | Cell Signaling Technology | 2541 | 1:500 |
| Human Phospho-Paxillin (Y31) Antibody, mouse | Invitrogen | 44720G | 1:1000 |
| Purified Mouse Anti-FAK, Clone 77/FAK | BD Transduction Laboratories | 610088 | 1:500 |
| Phospho-FAK (Tyr397) Recombinant Rabbit Monoclonal Antibody (31H5L17) | Invitrogen | 700255 | 1:500 |
| Akt (pan) (40D4) Mouse Monoclonal Antibody | Cell Signaling Technology | 2920 | 1:1000 |
| Phospho-Akt (Ser473) Antibody Rabbit | Cell Signaling Technology | 9271 | 1:500 |
| Glyceraldehyde-3-Phosphate Dehydrogenase (GAPDH) antibody mouse | Hytest | 5G4 / 5G4cc | 1:4000 |
| p44/42 MAPK (Erk1/2) (L34F12) Mouse Monoclonal Antibody | Cell Signaling Technology | 4696 | 1:1000 |
| Phospho-p44/42 MAPK (Erk1/2) (Thr202/Tyr204) (D13.14.4E) Rabbit Monoclonal Antibody | Cell Signaling Technology | 4370 | 1:1000 |
| Leupaxin Antibody Rabbit | Novus Biologicals | NBP2-55850 | 1:1000 |
| Src Antibody Rabbit | Cell Signaling Technology | 2108 | 1:1000 |
| Phospho-Src Family (Tyr416) Antibody Rabbit | Cell Signaling Technology | 2101 | 1:1000 |
| Anti-mouse IgG, HRP-linked Antibody | Cell Signaling Technology | 7076S | 1:5000 |
| Anti-rabbit IgG, HRP-linked Antibody | Cell Signaling Technology | 7074S | 1:5000 |
| Goat-anti-mouse 650 Azure secondary | Azure Biosystems | AC2166 | 1:2500 |
| Goat anti-rabbit secondary antibody AzureSpectra 800 | Azure Biosystems | AC2134 | 1:2500 |
| Goat-anti-mouse 800 Azure secondary | Azure Biosystems | AC2135 | 1:2500 |

### Fixed cells imaging

Fixed cells on gradient gels were imaged on a Nikon Eclipse Ti2-E with a 10x/0.3 CFI Plan Fluor objective (Nikon), using an Orca Flash 4.0 sCMOS camera (Hamamatsu Photonics) and 2 × 2 binning, or a Leica Thunder microscope with a 10x/ 0.45 HC Plan Apo objective (Leica), using a Leica K8 camera and 2 x 2 binning.

Fixed immunostained cells on homogenous hydrogels or glass-bottom dishes were imaged on a 3i Marianas spinning disk confocal microscope with a CSU-W1 scanning unit (Yokogawa), using a 40x/1.1 LD C-Apochromat objective (Nikon) and a Hamamatsu sCMOS Orca Flash4.0 camera.

To analyze cell density along the stiffness gradients, ImageJ was used to segment nuclei based on DAPI intensity thresholding and to quantify the beads intensity signal. Images were segmented in equal areas in which beads intensity and number of cells were quantified. Additionally, hexagonal maps of cell density based on their coordinates on the images were generated with R.

For morphological analysis of cells on homogenous gels, CellProfiler was used to mask the nucleus and the cytosol based on intensity thresholding of the DAPI and phalloidin signals, and to quantify YAP signal intensity within those masks. For the YAP cytosolic signal, a 3 µm ring around the nucleus within the cell mask was considered. ImageJ was used to manually segment individual cells, to measure their area and to segment focal adhesions based on automated intensity thresholding of the vinculin signal.

### Live cells imaging

Phase contrast live imaging was performed on a Nikon Eclipse Ti2-E with a 10x/0.3 CFI Plan Fluor objective (Nikon), using an Orca Flash 4.0 sCMOS camera (Hamamatsu Photonics). Images were recorded every 15min for 17h. The samples were kept at 37°C in a 5% CO_2_ atmosphere in complete growth medium supplemented with 12.5mM HEPES (pH7-7.5, Sigma-Aldrich) and 0.5mM sodium pyruvate (Sigma-Aldrich). The fluorescent beads indicating the stiffness gradients were imaged before and after live imaging recording.

Cells were segmented using ImageJ, based on manual intensity thresholding from the phase contrast images. Trackmate^62,63^ was used to automatically track cells and the migration metrics were extracted using a custom R script.

### Western Blot

To prepare samples for Western blot analysis, we plated cells in regular 6 well-plates or fibronectin-coated 0.5 and 25kPa SoftWell Easy Coat 6-well plates (Cell Guidance Systems) for 24h or the experimental time indicated in the figure. We then performed cell lysis in TXLB lysis buffer (1M Tris-HCl pH 7.5, 1M NaCl, 0.5% Triton-X100, 5% Glycerol, 1% SDS, 1X PhosStop phosphatase inhibitor (Sigma-Aldrich)) and scraped the cells. As the SoftWell plate wells absorb FBS from the medium which carries over to the lysates, a control well with complete medium without cells for each stiffness was used as a control. We boiled the samples at 90°C for 8 minutes and sonicated them for 10min. We quantified the protein concentrations using the DC protein assay kit from Biorad. The value obtained from the negative control well was subtracted to each sample from the same stiffness. We mixed equal amounts of proteins from each sample in sample buffer (80mM Tris-HCl pH6.8, 2.7% sodium dodecyl sulfate (SDS), 3.3% glycerol, 77mg/mL dithiothreitol (DTT), 0.5mg/mL bromphenol blue), boiled them at 90°C for 5 minutes and loaded them in Mini-Protean TGX Precast Gels 4-20% (BioRad). The samples were transferred on nitrocellulose membranes (BioRad) which were then blocked with AdvanBlock-Fluor blocking solution (Advansta) for 30min before incubation with primary antibodies diluted in TBS-0.1% Tween20 and AdvanBlock-Fluor blocking 1:1 solution overnight at 4°C. Fluorescent- or HRP-conjugated secondary antibodies were incubated in TBS-T TBS-0.1% Tween20 and AdvanBlock-Fluor blocking 1:1 1h at room temperature. The membranes were then imaged with Chemidoc (BioRad) or Azure Sapphire RGBNIR Biomolecular (Azure Biosystems) imagers.

Quantification of the bands intensity was done with ImageJ. To normalize each experiment, the intensity value of a band was divided by the average intensity of the same protein across all experimental conditions of the same replicate. Each value was then normalized by the corresponding GAPDH loading control.

### Mass cytometry

Cells where prepared for Masscytof analysis following the Maxpar surface staining protocol from Standard Biotools. Cells from one 10cm dish were detached using enzyme-free cell dissociation buffer (Gibco 13150016), washed and resuspended in serum-free medium. 1μM Cell-ID cisplatin (Standard Biotools, 201064) was added to the cells and incubated at room temperature for 5min. 5:1 Maxpar cell staining buffer (Standard Biotools, 201068) was added to quench the cisplatin and cells were spined down, washed and resuspended with cell staining buffer. Fc receptor blocking antibody (InVivoMAb anti-mouse CD16/CD32 Bioxcell, BE0307) was added (1:100) and incubated for 10min at room temperature. The antibody cocktail (see table 3) diluted in cell staining buffer was then added to obtained the desired antibody concentration and incubated for 30min at room temperature. Cells were washed twice in cell staining buffer before fixation with 1.6% formaldehyde (Pierce, 28908) for 10min at room temperature. Cells were then spined down and resuspended in Maxpar Fix and Perm buffer (Standard Biotools, 201067) with 500nM Cell-ID Intercalator-^103^Rh (Standard Biotools, 201103A) and incubated overnight at 4°C. The samples were then washed first with cell staining buffer and then with Maxpar water (Standard Biotools, 201069). They were finally resuspended in Maxpar water containing 1:10 EQ beads (Standard Biotools, 201078) and run in a Helios CyTOF (Standard biotools).

**Table 3:** Masscytof antibodies panel.

| Masscytof |  |  |  |  |
| --- | --- | --- | --- | --- |
| Antibody | Conjugated metal | Reference | Company | Dilution |
| CD41 | 89Y | 3089004B | Standard Bitools | 1:100 |
| CD325/N-Cadherin | 143Nd | 3143016B | Standard Bitools | 1:100 |
| CD138 | 145Nd | 3145003B | Standard Bitools | 1:100 |
| CD61 | 146Nd | 3146011B | Standard Bitools | 1:100 |
| HER2 (ErbB2/EGFR2) | 148Nd | 3148011A | Standard Bitools | 1:100 |
| CD34 | 149Sm | 3149013B | Standard Bitools | 1:100 |
| CD51/61 | 150Nd | 3150026B | Standard Bitools | 1:100 |
| CD102 (ICAM-2) | 151Eu | 3151015B | Standard Bitools | 1:100 |
| CD324 (E-Cadherin) | 158Gd | 3158018B | Standard Bitools | 1:100 |
| CD98 | 159Tb | 3159022B | Standard Bitools | 1:100 |
| CD49e | 160Gd | 3160015B | Standard Bitools | 1:100 |
| CD49b | 161Dy | 3161012B | Standard Bitools | 1:100 |
| Integrin b7 | 162Dy | 3162026B | Standard Bitools | 1:100 |
| CD49a | 163Dy | 3163015B | Standard Bitools | 1:100 |
| CD49F | 164Dy | 3164006B | Standard Bitools | 1:100 |
| Notch2 | 165Ho | 3165026B | Standard Biotoools | 1:100 |
| CD44 | 166Er | 3166001B | Standard Biotoools | 1:100 |
| Integrin a9b1 | 168Er | 3168013B | Standard Biotoools | 1:100 |
| CD24 | 169Tm | 3169004B | Standard Biotoools | 1:100 |
| CD54 | 170Er | 3170014B | Standard Biotoools | 1:100 |
| CD9 | 171Yb | 3171009B | Standard Biotoools | 1:400 |
| CD49d | 174Yb | 3174018B | Standard Biotoools | 1:100 |
| CD56 (NCAM) | 176Yb | 3176001B | Standard Biotoools | 1:100 |
| CD47 | 209Bi | 3209004B | Standard Biotoools | 1:100 |
| CD326 (EpCAM) | 141Pr | 3141006B | Standard Biotoools | 1:100 |
| CD151 (PETA-3) | 142Nd | 3142011B | Standard Biotoools | 1:100 |
| CD29 | 156Gd | 3156007B | Standard Biotoools | 1:100 |
| CD104 | 173Yb | 3173008B | Standard Biotoools | 1:100 |
| Purified anti-human EGFR Antibody Clone AY13 | 112Cd | #352902 | Biolegend | 1:50 |
| Purified anti-human CD10 Antibody Clone HI10a | 113Cd | #312202 | Biolegend | 1:50 |
| Purified anti-human CD166 Antibody Clone 3A6 | 147Sm | #343902 | Biolegend | 1:200 |
| Purified anti-human Notch 1 Antibody Clone MHN1-519 | 154Sm | #352102 | Biolegend | 1:50 |
| Purified anti-human Notch 3 (Maxpar® Ready) Antibody Clone MHN3-21 | 167Er | #345407 | Biolegend | 1:50 |
| Purified anti-human CD304 (Neuropilin-1) Antibody Clone 12C2 | 172Yb | #354502 | Biolegend | 1:50 |
| Human/Mouse Integrin alpha 11 Antibody | 106Cd | MAB4235 | R&D systems | 1:50 |
| Human ErbB3/Her3 Antibody | 110Cd | MAB3481 | R&D systems | 1:50 |
| Human Integrin alpha V/CD51 Antibody | 114Cd | MAB1219 | R&D systems | 1:50 |
| Human ErbB4/Her4 Antibody | 116Cd | MAB11311 | R&D systems | 1:50 |
| Human Syndecan-4 Antibody | 144Nd | MAB29181 | R&D systems | 1:50 |
| Human Integrin alpha V beta 5 Antibody | 152Sm | MAB2528 | R&D systems | 1:50 |
| Human Integrin beta 6 Antibody | 153Eu | MAB4155 | R&D systems | 1:50 |
| Human Integrin alpha 8 Antibody | 155Gd | MAB6194 | R&D systems | 1:50 |
| Human Integrin beta 8 Antibody | 175Lu | MAB4775 | R&D systems | 1:50 |
| CD49c a3 MAB1952z | 111Cd | MAB1952z | Sigma-Aldrich | 1:50 |

Data were analyzed on Cytobank (Beckman Coulter). All quantifications were made on live cells which were gated as cisplatin negative and Intercalator-^103^Rh positive.

### Single cell RNA sequencing sample preparation and processing

1.10^6^ cells were harvested for each sample. Samples were fixed with the Chromium Fixed RNA Kit, Human Transcriptome (10XGenomics). The cells were spined down, resuspended in fixation buffer (1X Fix & Perm buffer (10x Genomics) and 4% formaldehyde in nuclease-free water) and incubated overnight at 4°C. They were then spined down and resuspended in ice cold 1X quenching buffer (10x Genomics). 100µL of pre-warmed (65°C) enhancer and 275 µL of 50% glycerol were then added. Samples were kept at -80°C until further processing. Single cell libraries have been prepared at Single Cell Omics Core, Turku Bioscience Centre. Library preparation was performed according to the Chromium Fixed RNA Profiling protocol from 10XGenomics (CG000527) with the Dual Index Kit TS Set A (10XGenomics), using 600 000 cells as a starting amount and with a targeted cell recovery of 10 000 cells/sample. The quality of the samples was ensured using Agilent Bioanalyzer 2100. Sample concentration was measured with Qubit®/Quant-IT® Fluorometric Quantitation, Life Technologies. Sequencing was performed by Finnish Functional Genomics Centre – FFGC, Turku Bioscience Centre, University of Turku and Åbo Akademi University, Biocenter Finland infrastructure. Sequencing run was performed using Illumina NovaSeq 6000 S1 v1.5, with a targeted sequencing depth of 20 000 reads per cell and a read length of 28 + 90 bp.

### Single cell RNA sequencing analysis

#### Preprocessing and quality control

Base calling for single-cell RNA sequencing (scRNA-seq) was conducted with CellRanger (10X Genomics), and further processed using Seurat (v4). Filtered feature–barcode matrices were imported and merged across samples after appending sample-specific identifiers to retain unique barcodes. Features that were identified in less than 100 cells were excluded. Low-quality cells were removed by excluding cells with fewer than 500 detected genes or > 5 % mitochondrial reads. To discard doublets, samples were split by lineage and processed independently using DoubletFinder. The PCs that captured 90% of the cumulative variance were used as the number pf PCs for doubletFinder. For each sample, the expected doublet rate was estimated based on the number of recovered cells using canonical 10x Genomics multiplet rate estimates. Cells classified as singlets were retained for all downstream analyses. After duplicate removal, normalization and variance stabilization were conducted with SCTransform, regressing cell cycle markers (G2M & S phase). The first 10 PCs were used for UMAP visualization, and Louvain clustering was run with 0.1 resolution.

#### Trajectory and pseudotime-associated transcriptional shift

To explore continuous transcriptional trajectories, diffusion maps were computed with destiny R-package using PCA embeddings as input and cosine distance as the similarity metric separately for parental vs both metastatic lineages. Diffusion pseudotime was calculated to infer progression along the trajectory. To identify genes associated with the inferred transcriptional progression towards both metatstatic lineages, Spearman’s correlation coefficients (r) between genes and pseudotime were computed. These coefficients were used to establish genes associated with the transcriptional shift. Overlapping genes with absolute values of global r > 0.4 and parental lineage r > 0.2 were considered as significant. Gene set enrichment analysis was performed using fgsea with Gene Ontology Biological Process gene sets (MSigDB v2025.1). Genes were ranked by global pseudotime-associated correlation coefficients, and GSEA was conducted for the ranked gene list with fgsea R-package. Data visualization was conducted with ggplot2 and ComplexHeatmap R-packages.

### Bulk tumor analysis

Gene-expression data from the METABRIC-cohort was downloaded from cBioPortal (PMID: 22522925, PMID: 23550210). The cases were restricted to basal subtype, and a lymphovascular invasion (LVI) score was computed with the ssGSEA-algorithm utilizing genes previously shown to correlate positively with LVI^40^. LPXN mRNA expression was divided into quartiles, and statistical differences between these groups were assessed with Mann-Whitney U tests.

### Antibodies

#### Data representation and statistics

GraphPad prism 10.6.1 was used for most data graphical representation and statistical analysis unless when R was used as specified in the methods section. Three or more independent biological replicates were performed for experiments unless stated otherwise in the figure legends. Statistical tests are indicated in the figure legends. P values less than 0.05 were considered statistically significant and are provided in the figures, all P values are provided in the source data.

## Acknowledgements

We thank J. Siivonen and P. Laasola for technical assistance. We thank A. Isomursu, A. Taubenberger and the Ivaska laboratory for scientific discussions. For services, instrumentation and expertise, we would like to thank Cell Imaging and Cytometry Core (Turku Bioscience Centre, University of Turku) supported by Biocenter Finland, the Euro-BioImaging Finnish Node (Turku, Finland), Turku Bioscience Centre Single-Cell Omics Core Facility and the Finnish Functional Genomics Centre, both supported by the University of Turku, Åbo Akademi University and Biocenter Finland. This study has been supported by an ERC Advanced Grant (BorderControl; grant No. 101142305 to J.I.), the Finnish Cancer Institute (K. Albin Johansson Professorship; to J.I.), a Research Council of Finland Centre of Excellence (grant numbers 346131 and 364182, to J.I.), the Cancer Foundation Finland (J.I.), the Sigrid Juselius Foundation (J.I.), the Research Council of Finland’s Flagship InFLAMES (grant numbers 337530 and 357910) and the Jane and Aatos Erkko Foundation (J.I.). M.M. is supported by EMBO (ALTF 798-2021) and the Finnish Cancer Institute. M.V. is supported by ImmuDocs (National Doctoral Education Pilot Based on the Immune System). Funded by the European Union. Views and opinions expressed are however those of the author(s) only and do not necessarily reflect those of the European Union or the European Research Council Executive Agency. Neither the European Union nor the granting authority can be held responsible for them.

## Author contributions

Conceptualization: M.M., J.I. Methodology: M.M., J.I. Formal analysis: M.M., J.H. Investigation: M.M., J.H., M.V. Visualization: M.M., J.H., H.H. Writing: M.M., J.H., H.H., J.I. Supervision: J.I. Funding: J.I.

