## Extended Data for "Leupaxin inhibits the durotaxis and mechanosensitivity of metastatic breast cancer cells"

Extended Data Figures and Figure lengends:

Extended Data Figure 1, Related to Figure 1.

Extended Data Figure 2, Related to Figure 2.

Extended Data Figure 3, Related to Figure 2.

Extended Data Figure 4, Related to Figure 3.

Extended Data Figure 5, Related to Figure 4.

Extended Data Figure 6, Related to Figure 5.

Extended Data Figure 7, Related to Figure 6.

### Extended Data Figures

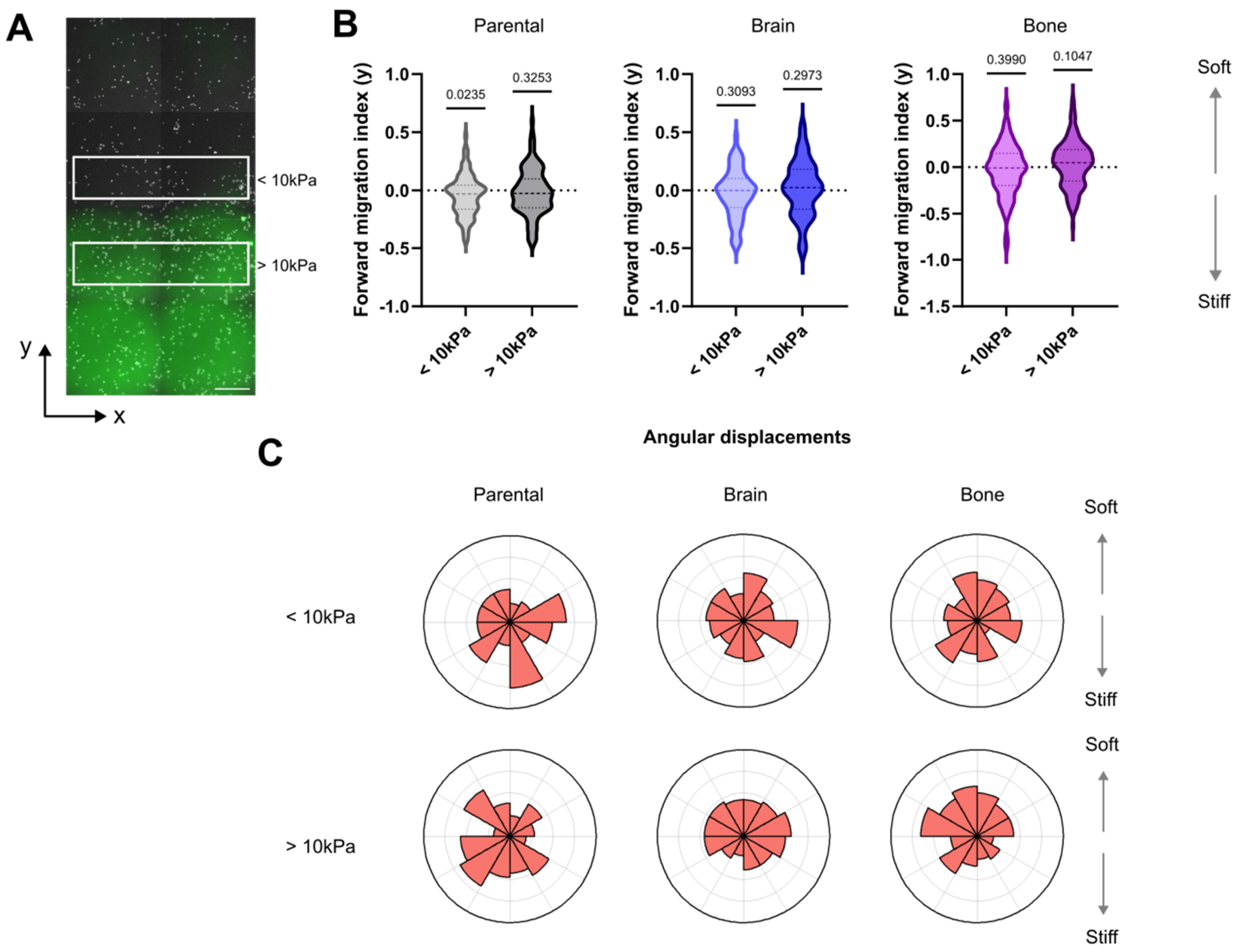


**Extended Data Figure 1:** MDA-MB-231 parental, brain and bone variants on stiffness gradient gels show different durotaxis phenotypes. **A)** A representative image of MDA-MB-231 parental cells during live imaging on a stiffness gradient gel. The bright-field image of the cells (grey) is overlaid with the fluorescence image of the beads (green). White boxes denote the soft and stiff regions of the gel used in cell tracking. Scale bar: 500µm. **B)** The forward migration index of tracked parental, -Br and -Bo cells along the stiffness gradient (y axis), on the soft (<10kPa) and the stiff (>10kPa) side. Median and quartiles are indicated as dashed lines on the violin plots. One sample Wilcoxon test. **C)** Angular displacements of the tracked cells. B,C) Parental: <10kPa N=94 cells, >10kPa N=107 cells; Brain: <10kPa N=129 cells, >10kPa N=156 cells; Bone: <10kPa N=118 cells, >10kPa N=132 cells; pooled from 3 independent experiments. Related to Figure 1.

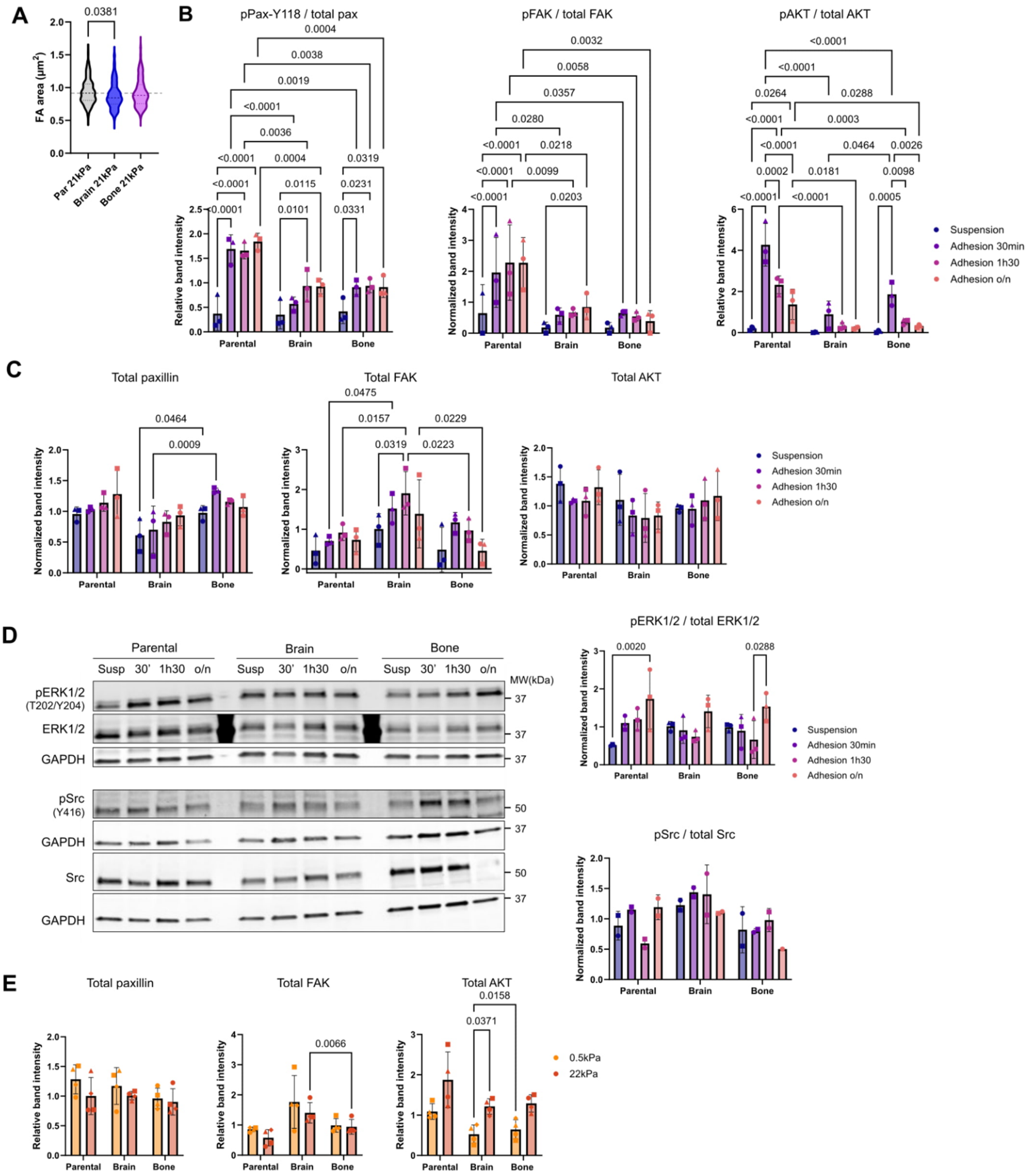


**Extended Data Figure 2:** Focal adhesion signaling in MDA-MB-231 parental, brain and bone variants. **A)** Mean FA area quantified from vinculin immunostaining in parental, -Br and -Bo cells on 22kPa gels. Parental N=134 cells; Brain N=130 cells; Bone N=106 cells, pooled from 3 independent experiments. Median and quartiles are indicated as dashed lines on the violin plots. Kruskal-Wallis test with Dunn's multiple comparisons test. **B,C)** Quantification from western blot analyses of the ratio of phosphorylated paxillin, FAK and AKT over the total corresponding proteins (B) and of total paxillin, FAK and AKT (C) comparing cells in suspension and following adhesion for 30 min, 1h30 and overnight (o/n). **D)** Representative western blots and corresponding analyses of the ratio of phosphorylated ERK1/2 and src over the total corresponding proteins comparing cells in suspension and following adhesion for 30 min, 1h30 and overnight (o/n). B,C,D) N=3 independent experiments (2 for pFAK in parental cells and for src). Mean ± SD. Mixed effects analysis with Tukey's multiple comparisons test. **E)** Quantification of western blot analyses of total paxillin, FAK and AKT in cells on soft (0.5kPa) and stiff (22kPa) gels. N=4 independent experiments. Mean ± SD. Two-way ANOVA with Tukey's multiple comparisons test. Related to Figure 2.

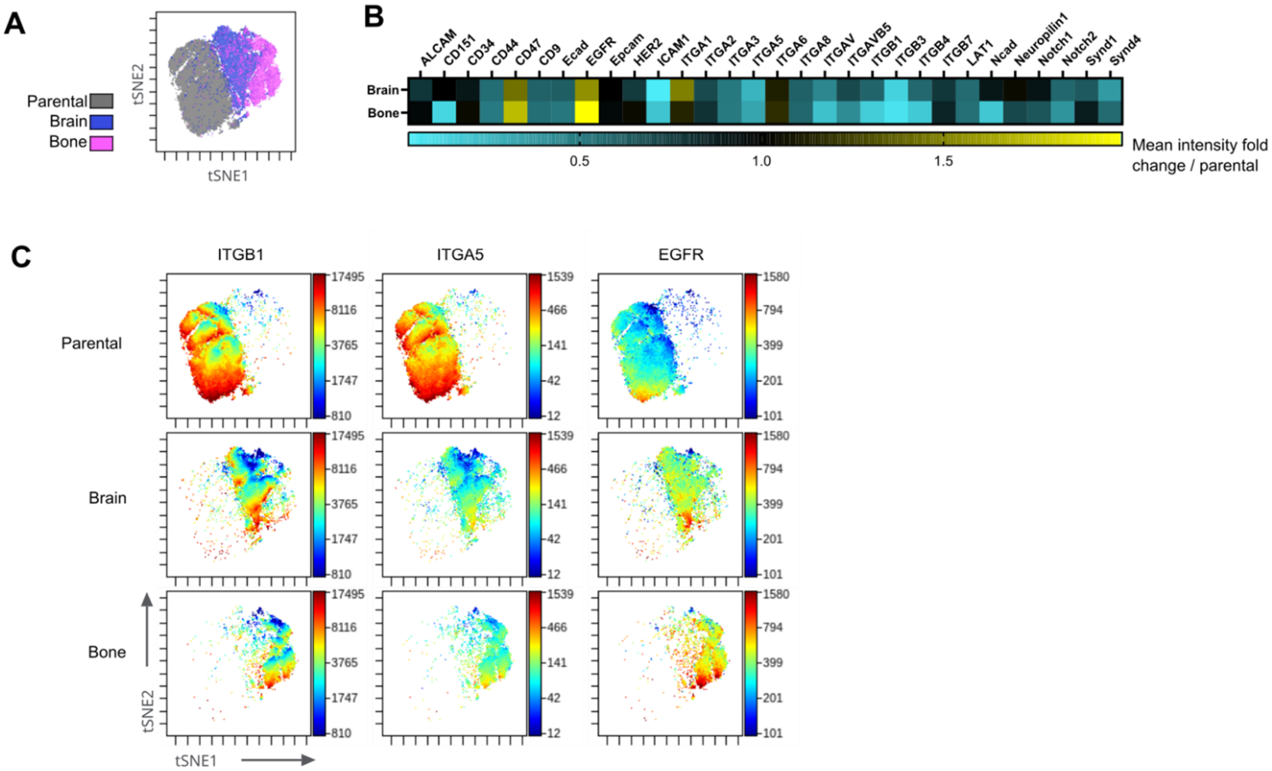


**Extended Data Figure 3:** MassCytof analysis of parental, -Br and -Bo cells. **A)** t-SNE analysis representation of data obtained from MassCytof analysis of the parental, -Br and -Bo cells. **B)** Heat map showing fold change expression of the indicated surface markers measured by MassCytof in -Br and -Bo cells compared to parental cells. **C)** Examples of surface marker expression in parental, -Br and -Bo cells from MassCytof analysis. A,B,C) Parental N=56,256 cells, Brain N=27,268 cells, Bone N=16,476 cells (after gating live cells). Related to Figure 2.

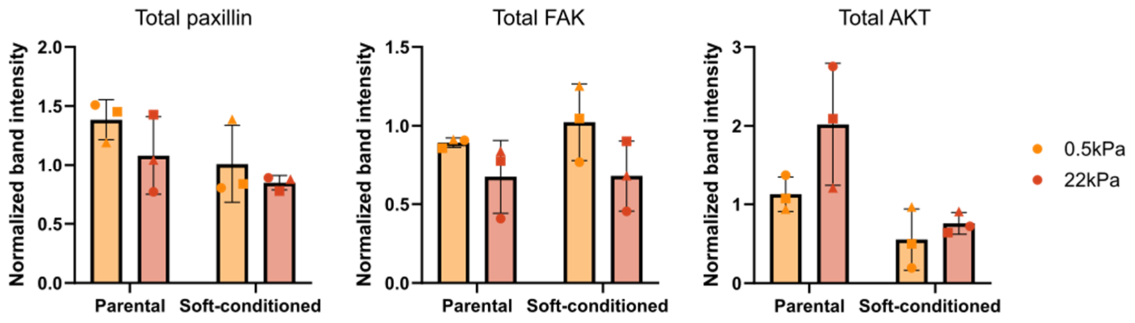


**Extended Data Figure 4:** Total paxillin, FAK and AKT levels are similar in MDA-MB-231 parental and soft-selected. Quantification of western blot analyses of total paxillin, FAK and AKT in cells on soft (0.5kPa) and stiff (22kPa) gels. N=3 independent experiments. Mean ± SD. Two-way ANOVA with Uncorrected Fisher's LSD. Related to Figure 3.

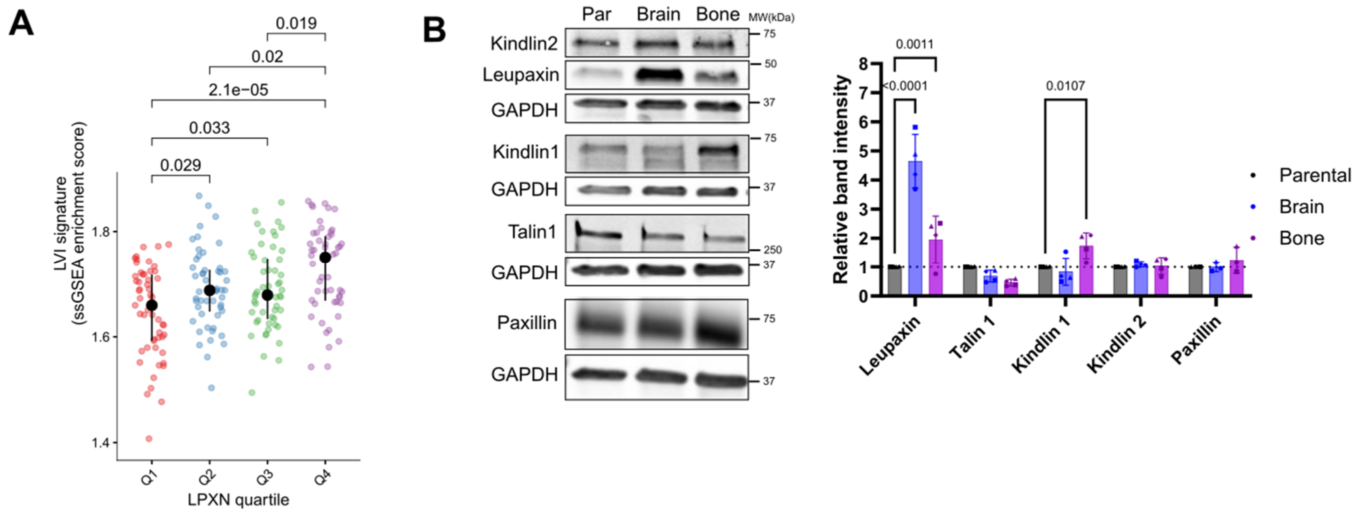


**Extended Data Figure 5:** Leupaxin clinical relevance and expression in cell lines compared to other FA proteins. **A)** Association between leupaxin mRNA expression from the breast cancer METABRIC-cohort restricting cases to the basal subtype (Q1-4: quartiles 1 to 4, lowest to highest expression) and lymphovascular invasion signature (LVI) in basal-subtype breast tumors (N = 209 patients). Mann-Whitney U test. **B)** Representative western blots and quantification of leupaxin, and the indicated FA proteins, in parental, -Br and -Bo cells. Mean ± SD. N=4 independent experiments for talin-1, kindlin-1 and -2 and leupaxin; N=3 independent experiments for paxillin. Two-way ANOVA with Dunnett's multiple comparisons test. Related to Figure 4.

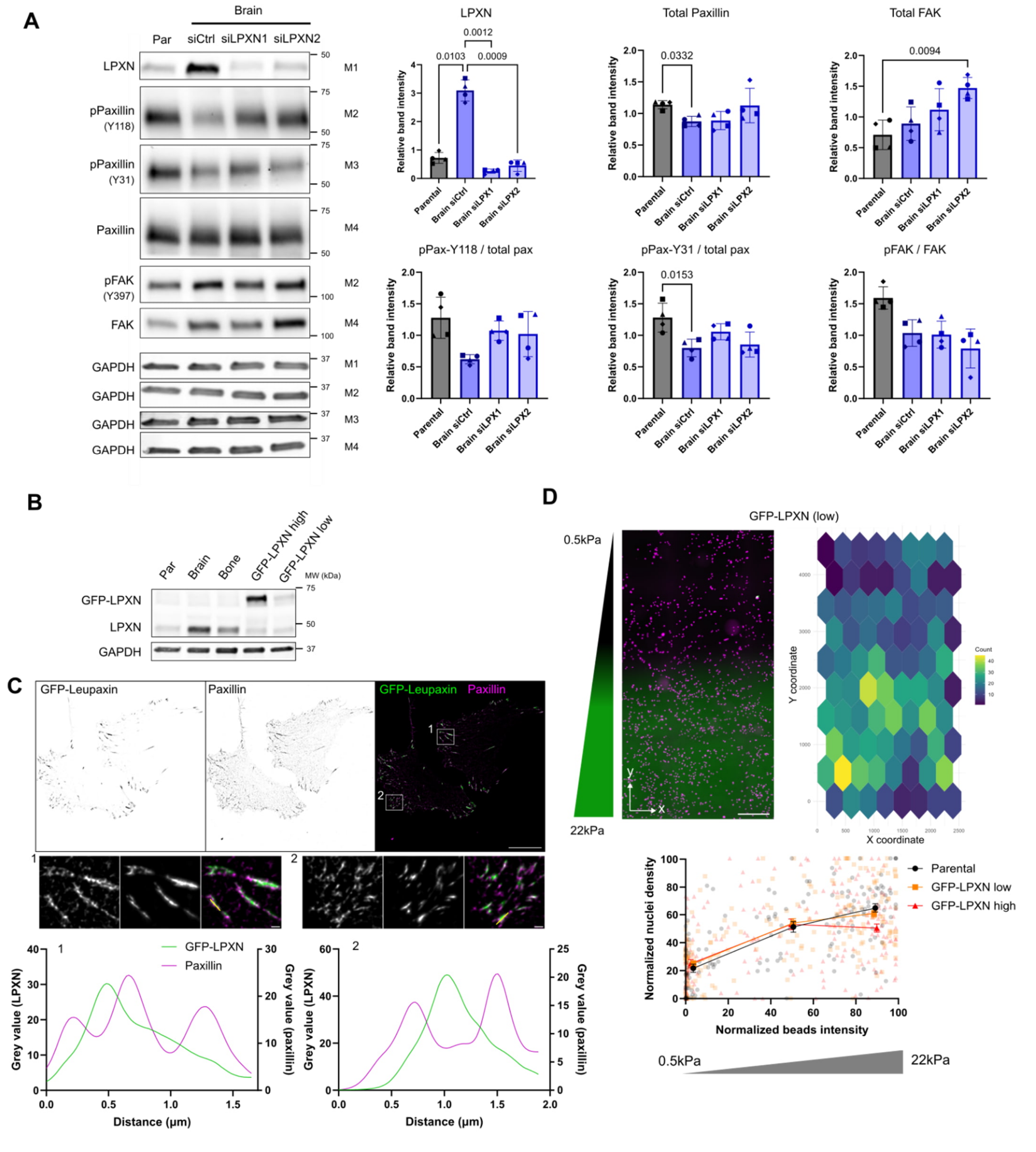

**Extended Data Figure 6:** Leupaxin silencing and overexpression. **A)** Representative western blots and quantification of the indicated proteins in parental cells and -Br cells transfected with a non-targeting siRNA (siCtl) or two different siRNAs targeting leupaxin (siLPXN1 and 2). N=4 independent experiments. Mean ± SD, RM one-way ANOVA with Tukey's multiple comparisons test for LPXN, Šídák's multiple comparisons test for pPaxillin-Y118 and -Y31, pFAK, total paxillin and FAK. **B)** Representative western blots showing endogenous LPXN and overexpressed GFP-LPXN in non-transfected parental, -Br and -Bo cells and in parental cells stably expressing high (GFP-LPXN high)

or low (GFP-LPXN low) levels of GFP-LPXN. **C)** Representative immunofluorescence images of paxillin and GFP-LPXN in GFP-LPXN high cells seeded on fibronectin-coated glass-bottom dishes. Image resolution was enhanced by eSRRF processing. Graphs indicate the paxillin and leupaxin intensity along the yellow lines indicated on the insets. Scale bar 20μm (full), 1μm (insets). **D)** A representative image of MDA-MB-231 GFP-LPXN low cells three days after seeding on a stiffness gradient gel. The corresponding heat maps of cell distribution and quantification of cell densities across the gradient are shown compared to parental and GFP-LPXN-high cells. Data were binned in three regions of interest corresponding to soft, intermediate and stiff areas of the gradients. Parental N=136 ROIs from 5 gels, GFP-LPXN high N=184 ROIs from 7 gels, GFP-LPXN low N=208 ROIs from 8 gels, pooled from 5 independent experiments, mean ± SEM, ordinary one-way ANOVA with Šídák's multiple comparisons test showed no significant difference between parental and GFP-LPXN low cells. Scale bar: 500μm. Related to Figure 5.

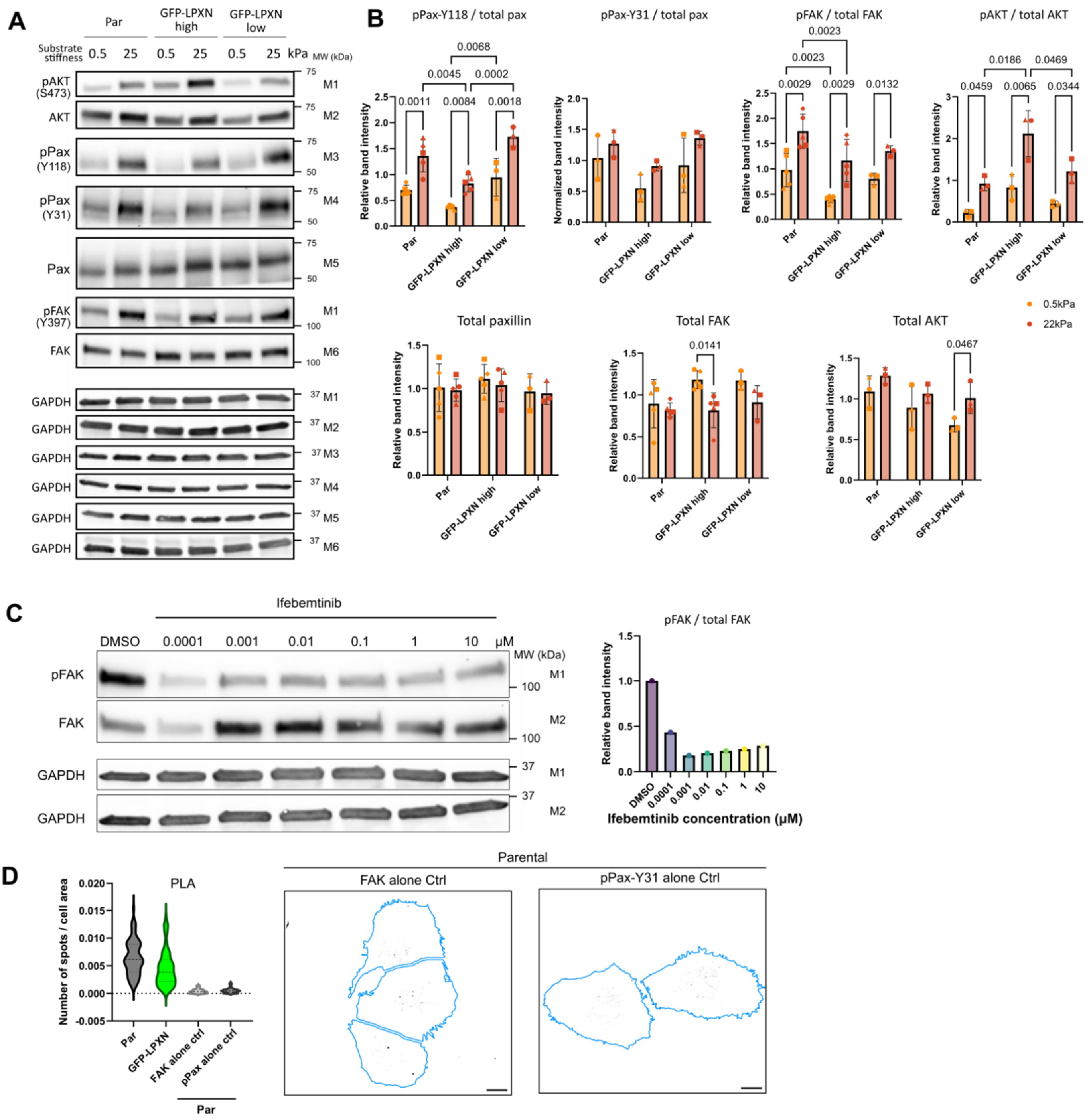

**Extended Data Figure 7:** Leupaxin overexpression and FAK inhibition. **A)** Representative western blots of parental, GFP-LPXN high and GFP-LPXN low cells after 24h on soft (0.5kPa) and stiff (25kPa) gels. **B)** Quantification of western blot analyses of the ratio of phosphorylated paxillin, FAK and AKT over the total corresponding proteins and of total paxillin, FAK and AKT in cells on soft and stiff gels. pPaxillin-Y118 and FAK / pFAK: N=5 independent experiments for parental and GFP-LPXN high, 3 independent experiments for GFP-LPXN low, mixed effects analysis with Tukey's multiple comparisons test. pPaxillin-Y31 and AKT / pAKT: N=3 independent experiments, two-way ANOVA with Tukey's multiple comparisons test. Mean ± SD. **C)** A western blot and corresponding quantification showing the effect of increasing doses of the FAK inhibitor, ifebemtinib, on FAK phosphorylation in MDA-MB-231 parental cells. **D)** Quantification of p-

Paxillin(Y31)–FAK PLA based on the number of spots in each cell divided by the corresponding cell area. Representative images of the controls using either p-paxillin(Y31) or FAK antibodies alone on parental cells are shown. Parental: N=77 cells, GFP-LPXN: N=90 cells, FAK alone: N=58 cells, p-paxillin alone N=55 cells, pooled from 3 independent experiments. Median and quartiles are indicated as dashed lines on the violin plots. Related to Figure 6.
